# High genetic diversity of bivalve transmissible neoplasia in the blue mussel *Mytilus trossulus* Gould from the subarctic Sea of Okhotsk

**DOI:** 10.1101/2021.12.07.471395

**Authors:** M.A. Skazina, N.A. Odintsova, M.A. Maiorova, L.T. Frolova, I.A. Dolganova, K.V. Regel, P.P. Strelkov

## Abstract

There are increasing findings of the bivalve transmissible neoplasia derived from the Pacific mussel *Mytilus trossulus (Mtr*BTN) in populations of different *Mytilus* species worldwide. The Subarctic is an area where this disease has not yet been sought despite the fact that *Mytilus* spp. are widespread there, and *M. trossulus* itself is a boreal species. We used cytological and histological techniques to diagnose disseminated neoplasia in a sample of *M. trossulus* from Magadan in the subarctic Sea of Okhotsk. Neoplasia was identified in 11 of 214 mussels studied. Using mtDNA COI sequencing, we revealed genotypes identical or nearly identical to known *MtrBTN* ones in the hemolymph of most of the diseased mussels. Both *Mtr*BTN evolutionary lineages have been identified, the widespread *Mtr*BTN2, and *Mtr*BTN1, so far only known from *M. trossulus* in British Columbia on the other side of the Pacific from Magadan. In addition, *Mtr*BTN2 was represented by two common diverged mtDNA haplolineages. These conclusions have been confirmed for selected cancerous mussels by molecular cloning of COI and additional nuclear and mtDNA genes. On the background of high genetic diversity, different cancers were similar in terms of ploidy (range 4.0 - 5.8n) and nuclear to cell ratio. Our study provides the first description of neoplasia and *Mtr*BTN in mussels from the Sea of Okhotsk and from the Subarctic, of both *Mtr*BTN1 and *Mtr*BTN2 in the same mussel population, and the first direct comparison between these transmissible cancers.

## Introduction

Bivalve molluscs, like most other metazoans, suffer from various cancers (Aktipis et al. 2015). A common cancer affecting bivalves is disseminated neoplasia (DN). In the course of this disease, neoplastic hemocytes, which are rounded and have enlarged pleomorphic nuclei and an unusual cytoskeleton, replace normal hemocytes and eventually infiltrate all the organs and tissues of the mollusc (Barber 2004; Carballal et al. 2015; Burioli et al. 2019; Odintsova 2020). DN may have a high prevalence and cause considerable damage to natural and commercial bivalve populations (Bower 1989; Farley et al. 1991; Elston et al. 1992; Villalba et al. 2001). It has recently transpired that DN can be a transmissible disease, passed from one mollusc to another by physical transfer of cancerous cells (Metzger et al. 2015, 2016). It is then referred to as bivalve transmissible neoplasia (BTN).

Blue mussels of *Mytilus edulis* species complex are widespread in temperate and subpolar seas of both Northern (*M. edulis, M. trossulus, M. galloprovincialis*) and Southern (*M. chilensis, M. platensis, M. planulatus, M. galloprovincialis*) Hemisphere (McDonald et al. 1991; Larraín et al. 2018). DN has been found in many geographical populations of mussels (see Ciocan and Sunila 2005; Wolowicz and Sokolowski 2006; Bramwell et al. 2021 for reviews and also below), and proven to be BTN in some cases (Metzger et al. 2016; Burioli et al. 2019; Yonemitsu et al. 2019; Hammel et al. 2021; Skazina et al. 2021), though mussels may have conventional (non-transmissible) DN as well (Hammel et al. 2021). Two evolutionary lineages of blue mussel BTN, *Mtr*BTN1 and *Mtr*BTN2, both derived from the Pacific mussel *M. trossulus*, have been recognized. *Mtr*BTN1 has been found only in *M. trossulus* from British Columbia (Northeast Pacific), while *Mtr*BTN2 has been reported from numerous populations of four different species of *M. edulis* species complex worldwide, including *M. trossulus* from the Sea of Japan (Northwest Pacific) (see Figure 1A for details). A widespread bipolar distribution of *Mtr*BTN2 indicates that it has spread together with infected mussels fouling ships (Yonemitsu et al. 2019; Skazina et al. 2021). *Mtr*BTN2 can be considered as a successful invasive species of a unicellular parasitic mussel, whose host range potentially encompasses the entire *M. edulis* species complex.

**Figure 1.**
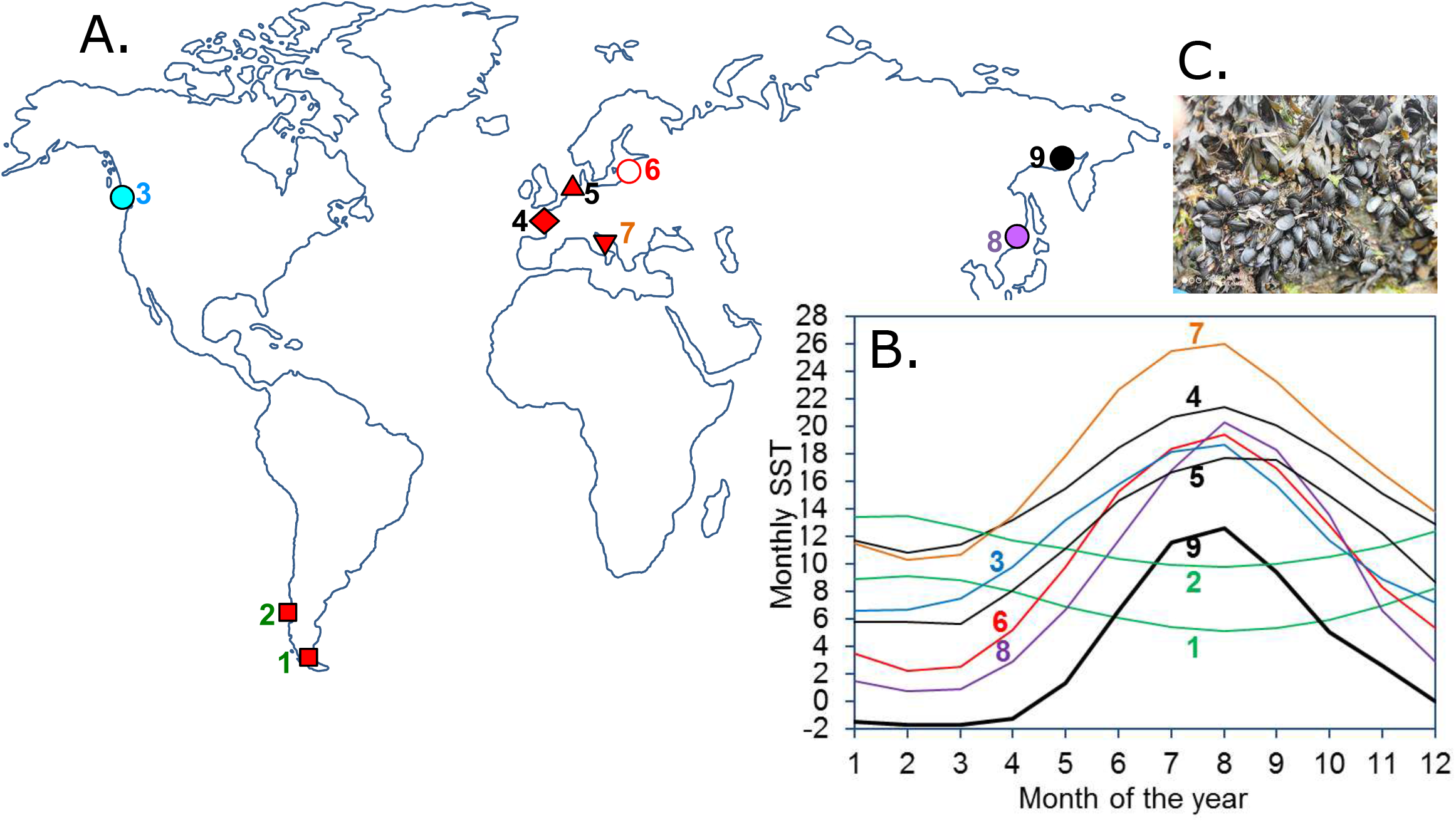
Geographical distribution of *Mtr*BTN and temperature conditions in the regions of its findings. (**A**) Geographical records of *Mtr*BTN. The colour of the symbols indicates the cancer lineage and the shape indicates the host mussel species. Blue – *Mtr*BTN1, red and violet – *Mtr*BTN2, black – *Mtr*BTN1 and *Mtr*BTN2 together, white – only mtDNA evidence of *Mtr*BTN2. Squares – *M. chilensis*, circles – *M. trossulus*, up-pointing triangle – *M. edulis*, down-pointing triangle – *M. galloprovincialis*, diamond – *M. edulis*, *M. galloprovincialis* and their hybrids. Numbers indicate places of *Mtr*BTN findings and data sources: 1 – Beagle Channel, Argentine (Yonemitsu et al. 2019), 2 – Chiloé Island, Chile (Yonemitsu et al. 2019), 3 – West Vancouver, British Columbia, Canada (Metzger et al. 2016), 4 – Atlantic coast from Arcachon Bay to Cherbourg Peninsula, France (Hammel et al. 2021), 5 – Wadden Sea, the Netherlands (Hammel et al. 2021), 6 – Gdansk Bay, the Baltic Sea, Poland (Zbawicka et al. 2014; Skazina et al. 2021), 7 – Istrian peninsula, the Adriatic Sea, Croatia (Hammel et al. 2021), 8 – Gulf of Peter the Great, the Sea of Japan, Russia (Skazina et al. 2021), 9 – Nagaev Bay, the Sea of Okhotsk, Russia (this study). (**B**) Monthly sea surface temperature (SST) in the areas where *Mtr*BTN was reported, after https://seatemperature.info/. Temperature curves for different areas are numbered as in plate A. Data from the following observation points closest to *Mtr*BTN findings were used: 1 – Ushuaia, 2 – Castro, 3 – West Vancouver, 4 – Arcachon, 5 – Harlingen, 6 – Sopot, 7 – Rovinj, 8 – Nakhodka, 9 – Magadan. (С) *Mtr*BTN-infected littoral mussels *M. trossulus* from Magadan. The largest mussels are about 50 mm in size.

The comparison of *Mtr*BTN1 and *Mtr*BTN2 is aggravated by their seemingly disjunct geographic distribution. In addition to the cancer genotype, the symptoms of this disease may depend on the genotype of the host, and the environment may play a role, too. A different degree of our knowledge of these two transmissible cancers is another aggravation. While *Mtr*BTN2 has been extensively studied cytologically and histologically (Burioli et al. 2019; Hammel et al. 2021; Skazina et al. 2021), *Mtr*BTN1 has not been examined in detail, though there are many older studies of DN in mussels from the Northeast Pacific (Farley 1969; Vassilenko and Baldwin 2014), where we now know this lineage occurs.

Since the discovery of *Mtr*BTN in 2016, six studies have been made, in which 48 cases of this disease have been reliably documented. Only seven of them (three *Mtr*BTN1 and four *Mtr*BTN2) have been reported from *M. trossulus*, the species in which *Mtr*BTN originated (Metzger et al. 2016; Riquet et al. 2017; Yonemitsu et al. 2019; Hammel et al. 2021; Skazina et al. 2021). This relative scarcity of studies is partly associated with a high labour intensity of the research.

Diagnosing *Mtr*BTN is a cumbersome procedure. To prove that an animal has transmissible cancer, it should be tested for genetic chimerism, and the genotype of the cancer should be distinguished from that of the host (Riquet et al. 2017). Potentially, rapid tests for *Mtr*BTN may be developed based on a parallel sequencing of the hemolymph, which is most affected by the disease, and the muscle tissue, which is least affected, at the COI mtDNA barcoding locus. Cancerous COI alleles do not seem to be very diverse (Yonemitsu et al. 2019; Hammel et al. 2021; Skazina et al. 2021) and have not been detected in healthy mussels, which means they should be relatively easy to identify. To note, in the four mussels with proven *Mtr*BTN2 from the Sea of Japan, host genotypes and cancer genotypes were successfully retrieved from, respectively, the foot muscle samples and the hemolymph samples by direct COI sequencing (Skazina et al. 2021).

BTN diversity and distribution in populations of its “parental” species *M. trossulus* and in blue mussel populations from high latitudes is an important question. *M. trossulus* is a boreal species of Pacific origin that has recently (post-glacially) invaded the North Atlantic (Rawson and Harper 2009; Wenne et al. 2016; Laakkonen et al. 2020). The distribution of *M. trossulus* (as well as *M. edulis*) extends northwards as far as latitudes 70-80 along different oceanic coasts (Golikov et al. 1998; Feder et al. 2003; Wenne et al. 2020). All the findings of BTN in *M. trossulus* were from the southern populations (regions 3, 6, 8 at Figure 1A), and most of its range remaining unexplored. So far, the northernmost report of *Mtr*BTN, (and DN) in blue mussels has been from *M. trossulus* in the Baltic Sea (Figure 1; Sunila 1987).

Our recent finding of BTN in *M. trossulus* in the Northwest Pacific (Skazina et al. 2021; Peter the Great Bay of the Sea of Japan; population 8 in Figure 1) set us thinking about the distribution and diversity of this disease along the Pacific coast of Siberia, which is discontinuously inhabited by *M. trossulus* (McDonald et al. 1991; Golikov et al. 1998). In this study, we looked for DN and BTN in *M. trossulus* mussels collected from the Sea of Okhotsk near Magadan. The annual sea surface temperatures in this subarctic sea are lower than in any other region where DN and BTN in blue mussels have ever been reported (Figure 1B).

## Materials and Methods

For the diagnosis of DN and BTN in mussels from the Sea of Okhotsk, we used the same stepwise approach as in our previous study of BTN in the Sea of Japan (Skazina et al. 2021), with some additions. The hemolymph was analysed by flow cytometry to quantify cell fractions of increased ploidy. Selected cancerous and healthy (control) mussels were also analysed using hemocytology in order to detail hemocyte morphology and tissue histology in order to check for infiltration of neoplastic hemocytes into mussel tissues and organs and to measure the hemocyte nuclear to cell ratio (NCR). To confirm BTN, the hemolymph and the foot tissues of four cancerous mussels were genotyped separately for three loci known to be diagnostic for *Mtr*BTN (Yonemitsu et al. 2019): nuclear elongation factor 1α gene (EF1α), mtDNA COI and control region (CR). Allele diversity was resolved by molecular cloning. The identity of the alleles with known *Mtr*BTN lineages was assessed by comparison with cancer genotypes from previous studies (summarized in: Yonemitsu et al. 2019; Skazina et al. 2021). The mussels for these analyses were selected on the basis of preliminary results on COI genotyping with standard LCO 1490 and HCO 2198 primers (Folmer et al. 1994). To assess the reliability of *Mtr*BTN identification by direct COI sequencing, we also sequenced COI from the hemolymph and the foot tissue of all 10 mussels with DN from our previous BTN study in the Sea of Japan (Skazina et al. 2021).

### Sample collection and preprocessing

Two samples of *Mytilus trossulus* were collected from the same littoral population at the mouth of Marchekan Creek in the Nagaev Bay within Magadan City (sampling coordinates 59°32’N; 150°46’E) in July 2020 (sample M1, sample size N=180) and September 2020 (sample M2, N=34). The molluscs were transported to the laboratory within 48 hours after sampling, and were kept there in the aquarium for a few days before the experiments. Fresh hemolymph was collected from the posterior adductor and visually checked for quality (the presence of hemocytes and the absence of contamination with gametes) under an inverted microscope CKX41 (Olympus, Japan). Aliquots of hemolymph were fixed for flow cytometry, hemocytology and genetic analyses according to the previously described protocols (Skazina et al. 2021). Small pieces of foot muscle tissues were fixed in 70% ethanol for genetic studies, while small pieces of mantle, gill and hepatopancreas were fixed for histological studies in 4% paraformaldehyde (PFA) in PBS, pH 7.8.

### DN diagnostics

Flow cytometry procedures were performed according to Vassilenko and Baldwin, 2014 as described in detail in Skazina et al. 2021. Mussels having at least 5% admixture of aneuploid cells among normal hemocytes in the hemolymph were classified as DN-suggested, while other mussels were classified as healthy. For all DN-suggested mussels and control healthy ones from sample M1 the diagnosis based on flow cytometry was verified by hemocytology and (or) histology.

The hemocytes were stained with TRITC-labelled phalloidin (Molecular Probes) and DAPI (Vector Laboratories) for visualization of the actin cytoskeleton and nucleus. We also used primary mouse monoclonal antibodies against anti-α-acetylated tubulin (clone 6-11B-1, Sigma Aldrich) for detecting mitotic spindles as described in Maiorova and Odintsova, 2015. Observations and images were made using the Zeiss LSM 780 confocal laser scanning microscope.

Histological sections 5 μm thick were stained with Meyer’s hematoxylin–eosin as described in Farley 1969 and Usheva and Frolova 2000 and examined using the light microscopes Primo Star or Axio Imager.Z2 (Zeiss, Germany) equipped with a digital camera. For each mussel, the cell and nucleus area (μm^2^) of at least five healthy and 30 neoplastic hemocytes randomly selected on slides was measured using AxioVision LE software (Carl Zeiss, Germany). NCR estimations were calculated as the ratios of nucleus and cell diameters recalculated from corresponding areas using formula 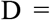 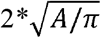, where D is diameter and A is area.

### BTN diagnostics and phylogenetic analysis

DNA extraction method, PCR primers for COI, CR and EF1α fragments, and cycling conditions were the same as in Yonemitsu et al. 2019. Preliminary diagnoses of cancerous mussels were done by direct Sanger sequencing of COI fragments amplified, separately, from the hemolymph and from the foot tissue. Primers LCO 1490 and HCO 2189 amplify the maternally inherited female mitochondrial genome (F-mtDNA) and, potentially, also the parentally inherited male mitochondrial genome (M-mtDNA) in mussels. Diverged M-mtDNA is detected only in the male gonads (Zouros 2013). Since we did not use the gonads, there was no risk of M-mtDNA amplification. Sequencing was performed using the Genetic Analyzer ABI Prism 3500xl in the Centre for Molecular and Cell Technologies of St. Petersburg State University Research Park (https://researchpark.spbu.ru/en/biomed-eng). PCR fragments of selected cancerous individuals were subjected to molecular cloning in Evrogen JSC facilities (Russia). The haplotype diversity of detected sequences was visualized as a TCS network by PopART software (Clement et al. 2002). Sequences detected only once or twice were considered as possible artefacts of the molecular cloning approach (Paabo et al. 1990; Bradley and Hillis 1997) and were excluded from further analyses. We also ignored variation in the polyA region of CR because polymerases may generate artefacts in mononucleotide regions (Clarke et al. 2001). Mussels studied by all three loci were used in the phylogenetic analysis. Maximum likelihood phylogenetic trees were created using parameters as in Yonemitsu et al. 2019 on the basis of MUSCLE alignments of newly obtained sequences from cancerous individuals along with collections of *Mtr*BTN-infected mussels studied by similar methods from Yonemitsu et al. 2019 and Skazina et al. 2021. Generation and visualization of alignments and trees were done in MEGA X software (Kumar et al. 2018).

In this study, we used allele nomenclature from Yonemitsu et al. 2019 and Skazina et al. 2021 for alleles detected in those studies. Newly discovered alleles were designated with the locus name and sequential number (e.g. COI-9), unless they were very similar to known cancer alleles, differing from them only by one or two substitutions. For such alleles, the original names were retained, and a sequential number was added (e.g. COI-U1 for an allele differing by one SNP from *Mtr*BTN1 specific COI-U).

### Analysis of the Sea of Japan collections

In our previous study of mussels from the Gaydamak population (sample J), 10 mussels were identified as cancerous by flow cytometry. Foot and hemolymph samples of four of these mussels plus the corresponding samples of two healthy ones were subjected to genotyping at COI, CR and EF1α and molecular cloning at CR and EF1α (Skazina et al. 2021). In this study we genotyped the remaining six cancerous mussels from the Gaydamak collection at COI using the procedures described above.

## Results

### DN diagnostics

Three distinct patterns were detected in the hemolymph of the studied mussels with the use of flow cytometry. (1) Most individuals had only one major diploid peak on the fluorescence histograms, sometimes with a minor admixture of tetraploid cells (<5%), and two cell types on the FSC-A/SSC-A dot plots, which were interpreted as normal hemocytes, granulocytes and agranulocytes (Figure 2A). These mussels were classified as healthy. (2) Nine individuals (M1-3, M1-25, M1-42, M1-96, M1-105, M1-111, M1-136, M1-162, M1-167) had an additional cell population, corresponding to aneuploid hemocytes with a higher DNA content (Figure 2C, D). The ploidy of these cells, calculated relatively to the 2n peak, ranged from 4.0 to 5.8 in different mussels and their proportion in hemolymph ranged from 15 to 97% (Table 1). (3) Two individuals (M1-95 and M2-22) had so few normal hemocytes (~1%) that the ploidy of aneuploid ones could not be estimated accurately (Figure 2B). Individuals with the second and the third pattern were classified as DN-suggested.

**Figure 2.**
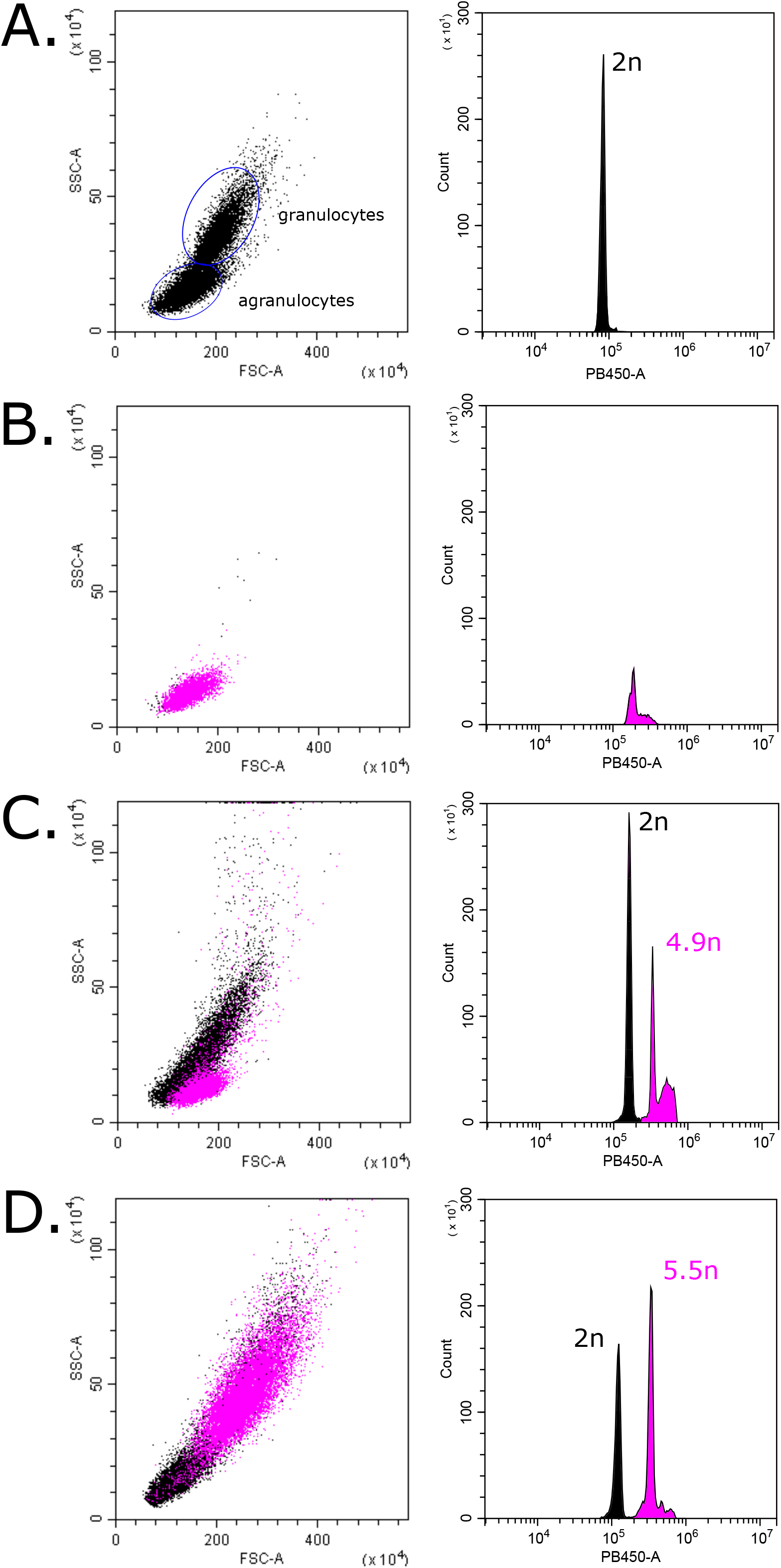
Flow cytometry patterns in the hemolymph of selected mussels from the Sea of Okhotsk. Each mussel is characterised by two diagrams. Dot plot on the left shows cell groups. The size of cells is on the OX axis (forward scatter, “FSC-A”) and their granularity on the OY axis (side scatter, “SSC-A”). Histogram on the right shows ploidy levels: the level of DAPI fluorescence is on the OX axis (“PB450-A”), the number of events is on the OY axis (“Count”). Relative ploidy levels are given near the peaks. In contrast to the healthy mussel (**A** – M1-12), an additional population of aneuploid cells (violet peaks and dots) is present in the hemolymph of cancerous mussels (**B** – M1-95 with *Mtr*BTN1 alleles, **C** – M1-96 cancerous mussel without *Mtr*BTN alleles, **D** – M1-162 with *Mtr*BTN2 alleles).

**Table 1.** Ploidy of aneuploid hemocytes, aneuploid cell rate and the COI-based BTN diagnosis of cancerous and healthy (control) mussels from the Sea of Okhotsk and the Sea of Japan. The relative ploidy of an aneuploid peak (“Ploidy”) is not given for mussels with deficiency of normal hemocytes. “Hemolymph genotype” and “Foot genotype” represent the COI genotypes detected by direct sequencing of DNA from the corresponding tissues. The mussels were classified into the following categories: “Healthy” (less than 5% aneuploid cells), “No BTN” (cancerous mussels lacking known *Mtr*BTN alleles and signs of genetic chimerism), “BTN1” and “BTN2” (cancerous mussels with alleles of the corresponding *Mtr*BTN lineage). Mussels, for which the diagnosis was confirmed by molecular cloning (Skazina et al. 2021; this study), are marked with asterisks.

| ID | Ploidy | Aneuploid cells rate | Hemolymph genotype | Foot genotype | BTN diagnosis |
| --- | --- | --- | --- | --- | --- |
| M2-22* | NA | 99% | COI-U1 | COI-U1 | BTN1 |
| M1-95* | NA | 99% | COI-U1 | COI-9 | BTN1 |
| M1-136 | 5.5 | 97% | COI-1 | COI-11 | BTN2 |
| M1-105* | 5.3 | 94% | COI-2 | COI-12 | BTN2 |
| M1-111 | 4.8 | 94% | COI-2 | COI-13 | BTN2 |
| M1-167 | 4.0 | 85% | COI-12 | COI-12 | No BTN |
| M1-162* | 5.5 | 57% | COI-1 | COI-8 | BTN2 |
| M1-96 | 4.9 | 45% | COI-14 | COI-14 | No BTN |
| M1-3 | 5.8 | 24% | COI-2 | COI-11 | BTN2 |
| M1-25 | 4.5 | 20% | COI-U1 | COI-11 | BTN1 |
| M1-42 | 5.8 | 15% | COI-1 | COI-12 | BTN2 |
| J1 | NA | 98% | COI-2 | COI-7 | BTN2 |
| J149 | 4.3 | 94% | COI-2 | COI-12 | BTN2 |
| J226 | 4.6 | 93% | COI-2 | COI-10 | BTN2 |
| J161* | 4.5 | 91% | COI-1 | COI-5 | BTN2 |
| J181* | 47 | 80% | COI-2 | COI-6 | BTN2 |
| J54* | 4.8 | 44% | COI-1 | COI-3 | BTN2 |
| J111* | 4.9 | 26% | COI-2 | COI-4 | BTN2 |
| J150 | 4.9 | 12% | COI-7 | COI-7 | No BTN |
| J162 | 5.2 | 5% | COI-4 | COI-4 | No BTN |
| J178 | 5.1 | 5% | COI-6 | COI-6 | No BTN |
| J17* | - | 0% | COI-7 | COI-7 | healthy |
| J38* | 4.0 | 3% | COI-8 | COI-8 | healthy |

Some differences in the hemolymph cell content were revealed between DN-suggested mussels (Supplementary Figure S1A-D) and control healthy ones (Supplementary Figure S1E, F) with the use of hemocytology. DN-suggested mussels had, in addition to normal hemocytes, also non-adherent hemocytes with enlarged pleomorphic nuclei and anomalous actin cytoskeleton. Tubulin staining did not reveal any mitotic activity in the hemolymph of any mussel.

Infiltration of neoplastic hemocytes with enlarged nuclei and basophilic cytoplasm into tissues was observed at the histological slides of the DN-suggested mussels (all DN-suggested mussels were examined in this respect except M2-22, M1-42 and M1-136) (Figure 3). The cell size and the nucleus size of healthy hemocytes were similar in different individuals. The characteristics of neoplastic hemocytes were also similar, with one exception. In M1-96, the size of the neoplastic hemocytes was noticeably smaller than in other individuals (by 23%), and so was the size of the nuclei (by 30%). The NCRs of neoplastic hemocytes were about twice as large as those of normal hemocytes (1.8 to 2.1 times larger in different mussels; the difference was the smallest in M1-96) (Table 2, Supplementary Figure S2). A few mitotic figures in neoplastic hemocytes were detected at all histological slides of the studied cancerous mussels.

**Figure 3.**
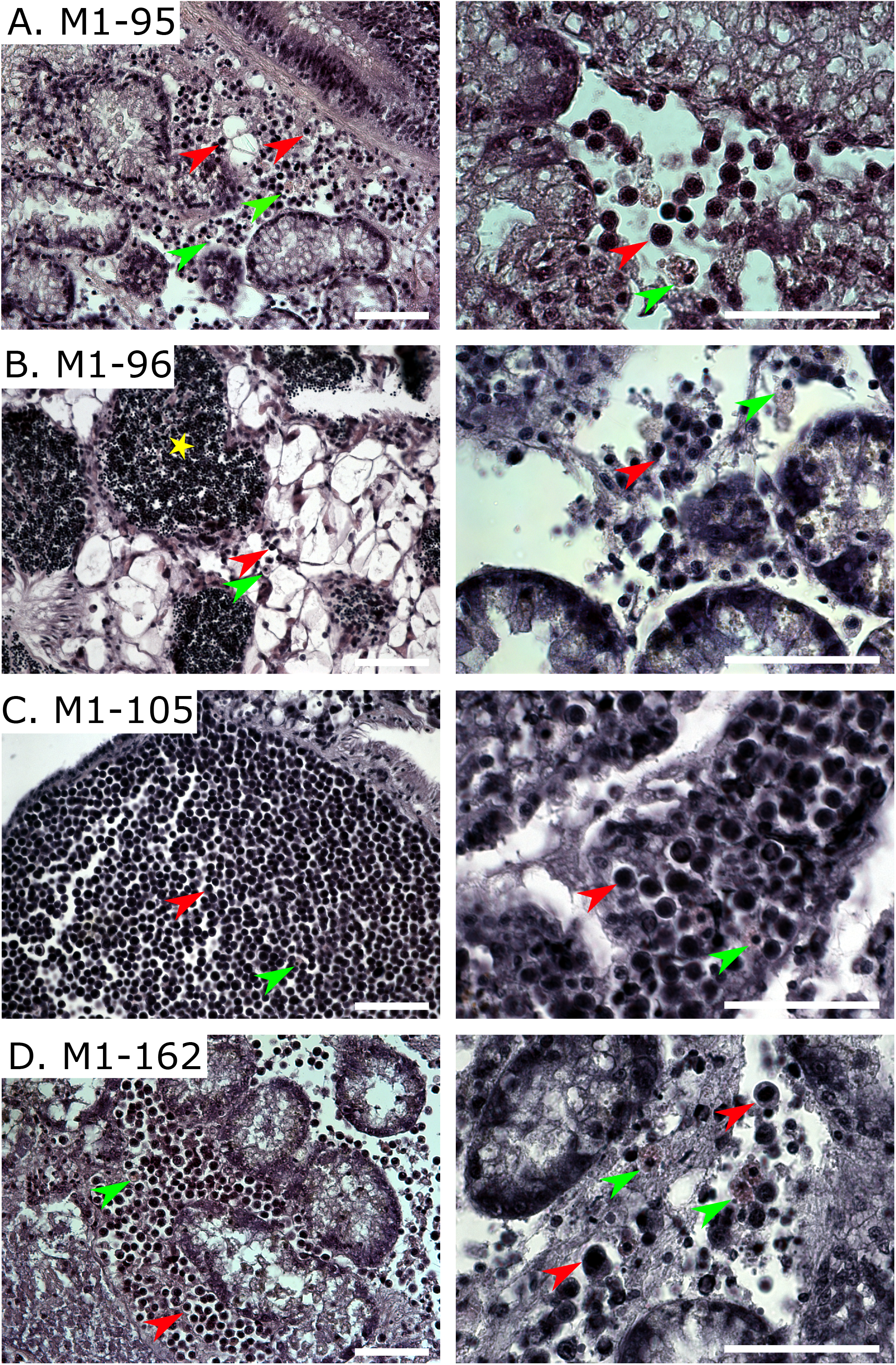
Histological sections of tissues of cancerous individuals stained with hematoxylin-eosin at 40x magnification (on the left) and 100x magnification (on the right). (**A)** Hemocyte infiltration in the connective tissues between digestive tubes of M1-95 individual infected with BTN1. (**B**) Hemocytes in the vesicular connective tissue of the gonad (on the left) and between digestive tubes (on the right) of M1-96 individual supposedly having conventional DN. (**C**) Heavy hemocyte infiltration of gill tissues of M1-105 individual infected by *Mtr*BTN2. (**D**) Infiltration of connective tissues between digestive tubes of M1-162 individual infected by *Mtr*BTN2. Red arrowheads point to cancerous hemocytes, green arrowheads point to healthy ones. A yellow asterisk marks the sperm cells. Scale bar, 50 μm.

**Table 2.** Morphometric characteristics of the neoplastic and the healthy hemocytes of cancerous mussels. Mean values, ranges and standard deviations (SD) of cell and nucleus diameter in μm and NCR are provided. For mussels with cancerous COI alleles *Mtr*BTN lineage is indicated in the left column.

| Individual | Cell type | N cells | Cell diameter | SD | Range | Nucleus diameter | SD | Range | NCR | SD | Range |
| --- | --- | --- | --- | --- | --- | --- | --- | --- | --- | --- | --- |
| M1-3<br>BTN2 | cancer | 30 | 6.6 | 0.55 | 5.66-7.91 | 5.18 | 0.53 | 4.38-6.52 | 0.78 | 0.05 | 0.7-0.88 |
|  | healthy | 5 | 8.22 | 1.5 | 6.54-10.53 | 3.15 | 0.22 | 2.85-3.42 | 0.39 | 0.08 | 0.31-0.52 |
| M1-25<br>BTN1 | cancer | 50 | 6.9 | 0.93 | 5.15-9.59 | 5.29 | 0.63 | 4.11-6.89 | 0.77 | 0.05 | 0.68-0.87 |
|  | healthy | 8 | 8.4 | 0.84 | 6.96-9.67 | 3.01 | 0.16 | 2.78-3.23 | 0.36 | 0.03 | 0.32-0.40 |
| M1-95<br>BTN1 | cancer | 50 | 6.58 | 0.64 | 5.26-7.64 | 5.05 | 0.54 | 4.03-5.9 | 0.77 | 0.04 | 0.66-0.87 |
|  | healthy | 16 | 7.91 | 0.9 | 6.84-10.71 | 2.98 | 0.29 | 2.57-3.4 | 0.38 | 0.05 | 0.24-0.44 |
| M1-96 | cancer | 30 | 5.4 | 0.68 | 4.34-6.73 | 3.73 | 0.44 | 3.14-4.48 | 0.69 | 0.09 | 0.51-0.81 |
|  | healthy | 6 | 7.89 | 1.30 | 6.38-9.65 | 3.05 | 0.27 | 2.61-3.36 | 0.39 | 0.07 | 0.32-0.49 |
| M1-105<br>BTN2 | cancer | 50 | 6.91 | 0.63 | 5.61-8.21 | 5.25 | 0.61 | 4.3-6.56 | 0.76 | 0.06 | 0.64-0.88 |
|  | healthy | 7 | 7.96 | 1.36 | 6.45-10.12 | 3.05 | 0.26 | 2.53-3.39 | 0.39 | 0.07 | 0.31-0.51 |
| M1-111<br>BTN2 | cancer | 50 | 6.89 | 0.77 | 5.77-8.36 | 5.07 | 0.66 | 4.09-6.5 | 0.74 | 0.06 | 0.55-0.83 |
|  | healthy | 7 | 7.77 | 0.89 | 5.97-8.45 | 2.96 | 0.4 | 2.39-3.49 | 0.38 | 0.06 | 0.3-0.48 |
| M1-162<br>BTN2 | cancer | 50 | 7.66 | 0.83 | 6.39-9.36 | 5.63 | 0.53 | 4.81-6.9 | 0.73 | 0.06 | 0.61-0.92 |
|  | healthy | 11 | 7.51 | 1.12 | 6.17-9.41 | 2.97 | 0.47 | 2.13-3.74 | 0.4 | 0.05 | 0.33-0.5 |
| M1-167 | cancer | 50 | 7.19 | 0.62 | 6.05-8.76 | 5.53 | 0.53 | 4.71-6.99 | 0.77 | 0.04 | 0.72-0.85 |
|  | healthy | 11 | 7.24 | 0.75 | 6.04-8.66 | 2.94 | 0.31 | 2.53-3.47 | 0.41 | 0.06 | 0.34-0.52 |

To sum up, DN was suggested for 11 mussels out of the 214 mussels studied by flow cytometry (5.1% of the sample). This diagnosis was confirmed by hemocytology and (or) histology for all cancerous mussels, except M2-22, which was not studied by these methods.

### BTN diagnostics

#### Direct COI sequencing

Results of the COI genotyping of the hemolymph and the foot tissues of 11 cancerous mussels from Magadan, 10 cancerous mussels and two healthy control mussels from the Sea of Japan are provided in Table 1, 2 and Supplementary Figure S3. Visual inspection of the chromatograms revealed the following three patterns. (1) Distinct sequences in different tissues (M1-95, M1-105, M1-111, M1-136, J1, J149, J226). (2) Overlapping peaks (“piggybacks”) in multiple positions on chromatograms from the hemolymph, from the foot or from both tissues (M1-3, M1-25, M1-42, M1-162, J54, J111, J161, J181) (Supplementary Figure S3). This pattern was interpreted as a mixture of two different sequences, a cancerous one with more pronounced peaks in the hemolymph than in the foot and a host one with more pronounced peaks in the foot tissues (see Supplementary Figure S3 capture for more explanations). (3) Identical sequences in both tissues with no signs of heteroplasmy (the rest of mussels). In terms of the rate of aneuploid cells, all mussels with the first pattern were severely diseased (>93% of aneuploid cells), while those with the second pattern were moderately to severely diseased (15-91%). The third group was very heterogeneous and included a severely diseased M2-22 (99%), moderately diseased M1-167 and M1-96 (85% and 45% respectively), slightly diseased J150, J162 and J178 (5-12%) and control healthy mussels J17 and J38 (Table 1).

The comparison of the nucleotide sequences detected in newly studied mussels (not considering six previously genotyped mussels from J sample) with the collection of known *Mtr*BTN genotypes revealed alleles characteristic of *Mtr*BTN2 (COI-1 or COI-2) in the hemolymph of nine mussels including six from Magadan. Further, a COI-U1 allele differing by 1 SNP from the unique U allele of *Mtr*BTN1 lineage was revealed in the hemolymph of M1-95, M1-25 and M2-22. The same COI-U1 allele was detected in a homoplasmic state in the foot of the heavily infected M2-22 individual. The other alleles identified in our study differed by 2-12 substitutions from the known *Mtr*BTN alleles and by 3-11 substitutions from each other. They were most probably the alleles of the host. No known cancer alleles and no signs of genetic chimerism were revealed in M1-96 and M1-167 individuals, which had a moderate rate of aneuploid cells, and in J150, J162 and J178 individuals, which has a low rate of aneuploid cells <15% (Table 1).

Ploidy and mean NCR of neoplastic hemocytes from mussels collected near Magadan marked by different BTN COI genotypes were as follows: *Mtr*BTN2 with COI-1 (ploidy range 5.5-5.8n; NCR 0.73), *Mtr*BTN2 with COI-2 (4.8-5.8n; NCR range 0.74-0.78), *Mtr*BTN1 with COI-U1 (4.5n; 0.77), M1-96 (4.9n; 0.69), M1-167 (4.0n; 0.77) (Table 1, 2; Supplementary Figure S2). Ploidy of neoplastic hemocytes in mussels with *Mtr*BTN2 alleles from the Sea of Japan (range 4.3-4.9n) was on average lower than that in mussels from the Sea of Okhotsk (4.8-5.8n) (Table 1). These differences were statistically significant at p=0.05 (the two-sample Kolmogorov-Smirnov test calculated in PAST; Hammer et al. 2001). In a similar comparison between all cancers from two samples, the differences were insignificant.

Four mussels were selected for detailed analysis by molecular cloning. They were M1-95 and M2-22, which presumably had *Mtr*BTN1, and M1-105 and M1-162, which presumably had *Mtr*BTN2 with different COI alleles.

Results of molecular cloning are summarized in Supplementary Table S1 (list of all sequences detected), Supplementary Figure S4 (TCS haplotype network of all sequences) and Table 3 (frequencies of common alleles). Nucleotide sequences of alleles found in this study are available at NCBI GenBank (Accession numbers MZ724668-MZ724679 for COI, MZ751085-MZ751090 for CR and MZ751091-MZ751101 for EF1α, Supplementary Table S2).

**Table 3.**
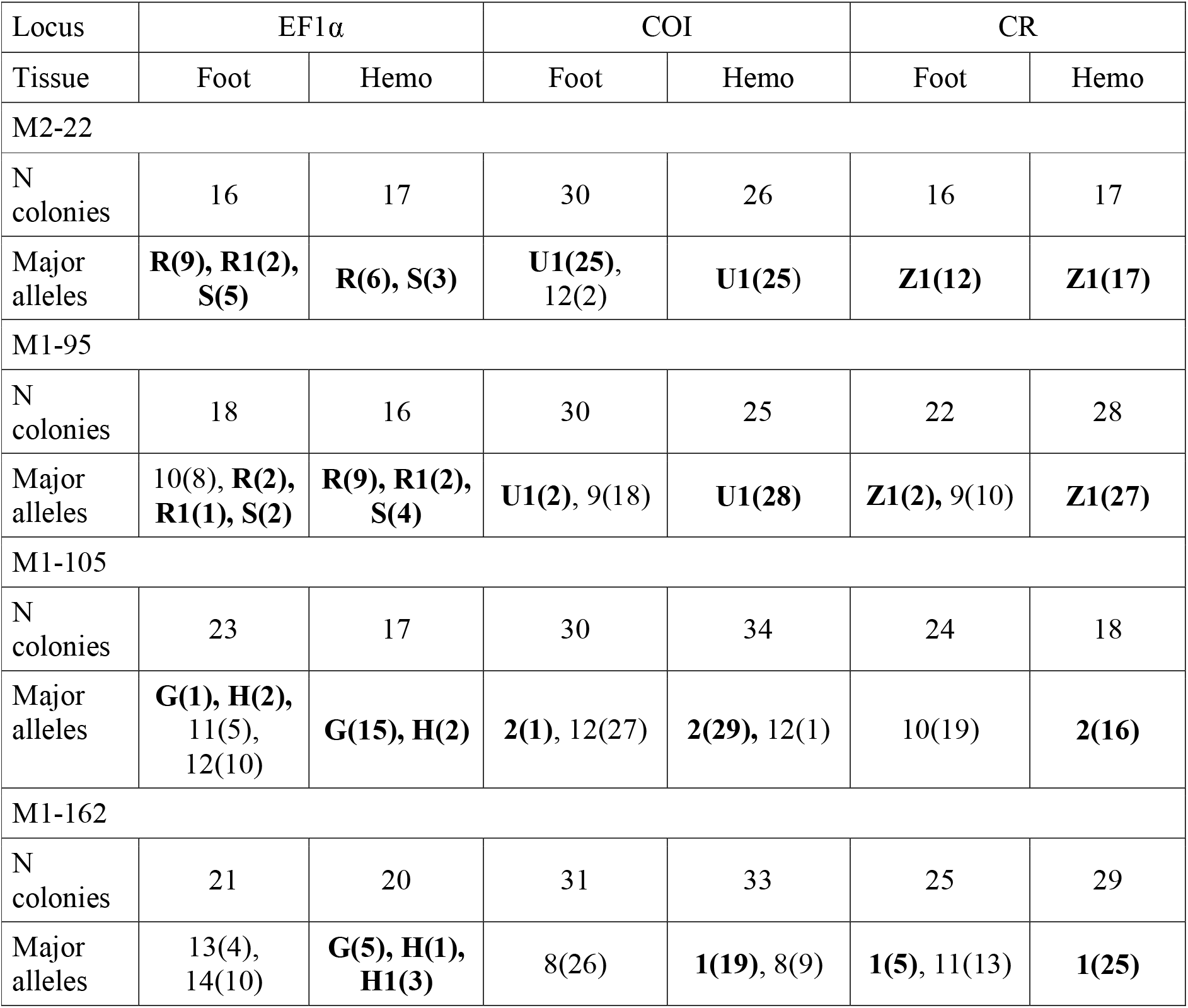
Sequences detected by molecular cloning in this study. The number of colonies with corresponding sequence is given in brackets. Sequence names shown in bold corresponds to cancerous alleles.

Cloning confirmed the preliminary diagnosis based on direct COI sequencing. At EF1α, M1-105 and M1-162 bore known combinations of unique *Mtr*BTN2 alleles (G and H) while M2-22 and M95 bore known combinations of unique *Mtr*BTN1 alleles (R and S). At mitochondrial loci, different characteristic *Mtr*BTN2 haplotypes were identified in M1-162 (COI-1, CR-1) and M1-105 (COI-2, CR-2). M2-22 and M1-95 were marked by a combination of COI-U1 and CR-Z1 alleles, CR-Z1 differing by two substitutions from the characteristic CR-Z of BTN1. (To remind, we do not consider here the variation at the polyA region of CR as putatively artefactual; information on this issue is provided in the Supplementary Table S3). M2-22, M1-95 and M1-162 were seemingly trisomic at EF1α due to the “additional”, relatively rare R1 (M2-22, M95) or H1 (M1-162) alleles, differing by one substitution from the common R and H alleles, respectively (Table 3). This extra variation could be explained by methodical artefacts of cloning or by somatic mutations in aneuploid cancers. Aneuploidy also could explain the misbalance between common cancerous EF1α alleles: R allele was represented by more clones than S allele in *Mtr*BTN1 (noted also by Metzger et al. 2016) and G allele was represented by more clones than H allele in *Mtr*BTN2 (Supplementary Figure S4). Other alleles identified were unrelated to cancerous ones and were usually more common in the foot tissues than in the hemolymph (Supplementary Figure S4; Table 3). These alleles evidently represented mussel host genotypes.

Salient features of transmissible cancers of mussels from the Sea of Okhotsk against the background of their global diversity can be seen in phylogenies in Figure 4. *Mtr*BTN2 mtDNA genotypes from the Sea of Okhotsk were nearly identical to those from the Sea of Japan. In particular, both samples were polymorphic for two diverged haplolineages marked by combinations of COI-1+CR-1 and COI-2+CR-2 alleles. These Northwest Pacific populations were different from European, Argentinian and Chilean ones. The macro-geographical variability of *Mtr*BTN2 has been discussed in detail in Yonemitsu et al. 2019 and in Skazina et al. 2021. In brief, two distinct CR alleles similar to CR-1 and CR-2 from the Northwest Pacific (CR-D and CR-C by classification of Yonemitsu et al. 2019, respectively) were observed elsewhere, in presumably heteroplasmic (Argentine, both alleles detected in the same cancerous mussels) or homoplasmic (CR-C in Chile and CR-D in Europe) states. COI diversity in Europe and Argentina was considerably lower than in the Northwest Pacific. Chilean *Mtr*BTN2 samples were the most distinctive because their COI alleles were more related to *M. chilensis* than to *M. trossulus*. *Mtr*BTN1 genotypes from Magadan differed from those from the British Columbia ones by 1-2 substitutions at mitochondrial loci and an “additional” R1 allele at EF1α.

**Figure 4.**
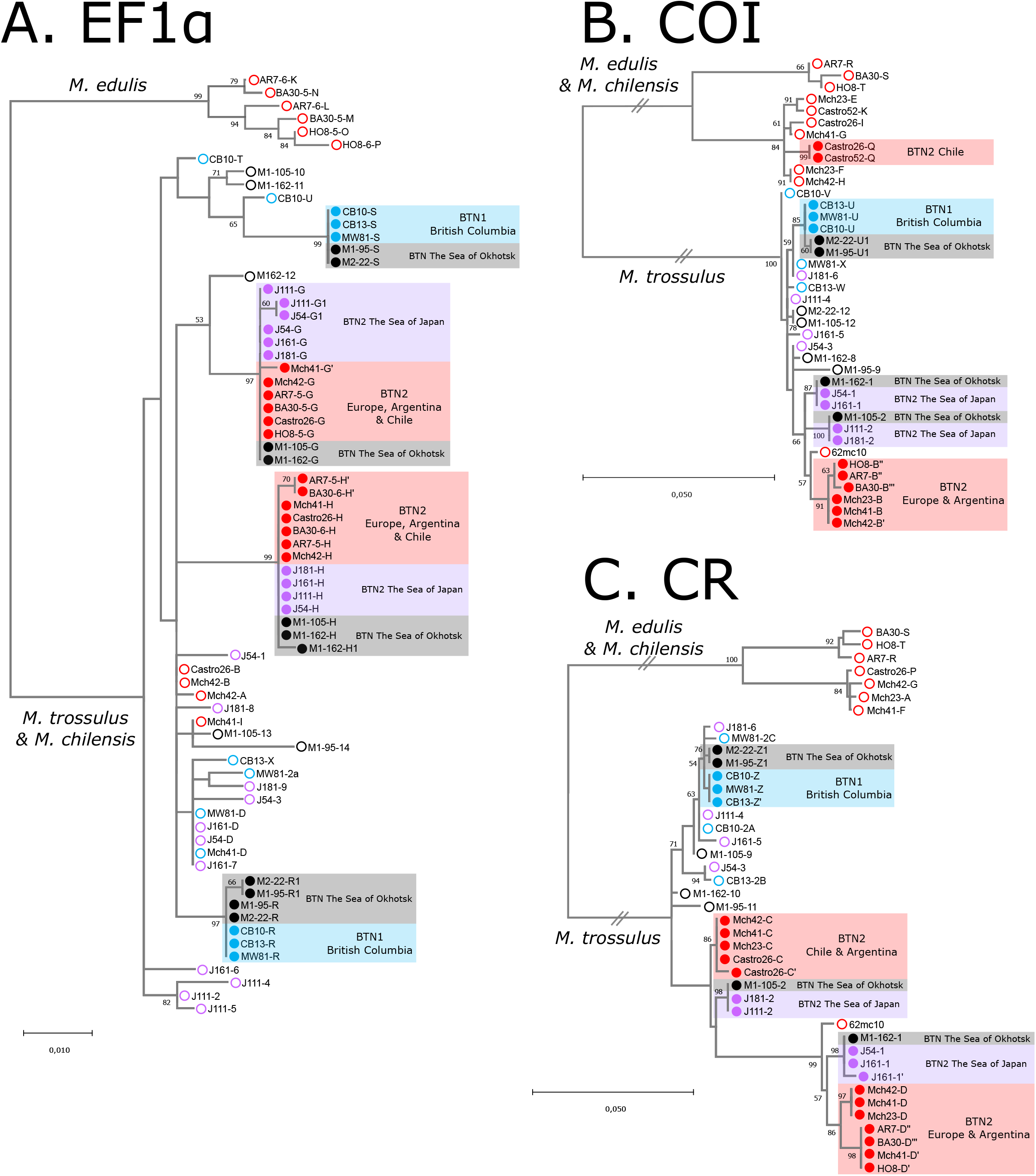
Phylogenetic analysis of nuclear and mitochondrial loci of mussels with BTN. Maximum likelihood (ML) trees are rooted at the midpoint, bootstrap values below 50 are removed. (**A**) EF1α ML tree based on the 630 bp alignment (HKY+G). (**B**) COI ML tree based on the 630 bp alignment (GTR+G). (**C**) CR ML tree based on the 828 bp alignment (HKY+G). Branches marked by two slashes are shortened 5-fold for COI and CR. The scale bars mark genetic distance. The analysis is based on 10 diseased mussels from Yonemitsu et al. 2019 (marked by red colour; note that Castro26 host COI allele J was not included because of the short sequence length), one mitochondrial sequence from the Baltic mussel 62mc10 (Zbawicka et al. 2014, also marked by red colour), four diseased mussels from the Sea of Japan (Skazina et al., 2021; marked by violet colour), and new data on four mussels with DN from the Sea of Okhotsk (marked by black colour). Open circles at the end of branches mark host alleles, closed circles mark cancer alleles. Geographic origin of the samples and species identity of the clades are indicated.

## Discussion

In this study we showed the presence of DN in mussels *Mytilus trossulus* from the subarctic Sea of Okhotsk. In some cases, this disease was represented by BTN. Though our sample was small, both known evolutionary lineages of BTN, *Mtr*BTN1 and *Mtr*BTN2, were found in it. While *Mtr*BTN2 is a widespread lineage, *Mtr*BTN1 has previously been described only in mussels from Western Canada, about 5000 km across the Pacific from our collection site near Magadan (Figure 1A). This is the first report of *Mtr*BTN1 and *Mtr*BTN2 from the same mussel population. Below, we discuss the patterns of geographical distribution of *Mtr*BTN, the genetic features of its Northwest Pacific clones, the possibilities of developing rapid tests for BTN based on direct COI sequencing as well as cytological differences and similarities between *Mtr*BTN1 and *Mtr*BTN2.

### Geographical and climatic limits of MtrBTN distribution

Our finding of transmissible cancer in mussels collected near Magadan extended the known distribution of this disease into the Pacific Subarctic. It is too early to judge about its overall geographical distribution because many regions such as the eastern coast of North America and Northern Europe have not been surveyed. Nevertheless, the available data on the geographical and temperature boundaries of *Mtr*BTN, which are summarized in Figure 1, allow us to draw some preliminary conclusions about the temperature preferences at least of *Mtr*BTN2, which is much more widespread than *Mtr*BTN1.

Similarly to its hosts, the blue mussels themselves, *Mtr*BTN2 survives across almost the entire temperature range of temperate seas, including subpolar seas with strong (Magadan, in *M. trossulus*, region 9 in Figure 1) and weak (Beagle Strait, in *M. chilensis*, region 1 in Figure 1) seasonality. In the warmest place where it has been found (Adriatic, in *M. galloprovincialis*, region 7 in Figure 1), it tolerates temperature regimes unacceptable for its parental species, *M. trossulus*, which is boreal and requires temperatures below 17°C to maintain a positive scope for growth (Fly and Hilbish 2013). In the Adriatic Sea, the sea surface temperatures (SST) are above this limit for most of the year (Figure 1B). How far northwards *Mtr*BTN spreads and whether it is present in the Arctic remain an open question. Based on the available data, it cannot be ruled out that the spread of the deadly *Mtr*BTN is not limited by either climate or geography and that any mussel population is potentially at risk.

### Genetic features of MtrBTN in the Northwest Pacific

The material on *Mtr*BTN1 is very limited, being represented by two tiny samples from the opposite coasts of the Northeastern Pacific (three cancers from Canada studied by Metzger et al. 2016, and two from Russia, Figure 4). Based on this material, we can only acknowledge minor qualitative differences between these cancers, namely, that they differ by one substitution at COI fragment (0.16% net divergence), by two substitutions at CR fragment (0.32%) and by the presence of an “additional” EF1α-R1 allele in cancers from Magadan. However, this apparent trisomia could be an artefact of the genotyping method.

The spectrum of *Mtr*BTN2 genotypes from the Sea of Okhotsk (this study) was the same as in the Sea of Japan (Figure 4). It is likely that the cancers from these two seas represent the same broad geographic population. If so, there is little to add to the discussion of macrogeographic variability of *Mtr*BTN2 provided in our previous paper (Skazina et al. 2021). One noteworthy fact is that *Mtr*BTN2 from the Northwest Pacific, on the one hand, and *Mtr*BTN2 from invasive populations of Argentina and Western Europe, on the other hand, show relatively high levels of COI net divergence (0.95-1.27% *vs* 0.1-0.3% between Argentina and Europe, Figure 4). This observation may imply that the Northwest Pacific was not the direct source of the invasion. Other possible explanations are a long history of invasive populations or a high mutability of mtDNA.

Based on our data, we may now answer the question whether Northwest Pacific *Mtr*BTN2 is homoplasmic or heteroplasmic by two diverged haplolineages marked by combinations of COI-1+CR-1 and COI-2+CR-2 alleles. Yonemitsu et al. 2019 identified both diverged CR alleles (similar to CR-1 and CR-2) in all the three cancerous mussels from Argentina studied by molecular cloning and showed using qPCR that the levels of two alleles could be very different. Based on these results, it was suggested that the homoplasmy by these alleles observed in other populations, also in the Sea of Japan, may be a seeming one, the minor allele eluding molecular cloning (Yonemitsu et al. 2019; Skazina et al. 2021).

In this study we sequenced numerous colonies with CR and COI from two mussels with *Mtr*BTN2 from Magadan (in total 416) and four mussels from the Sea of Japan involved in our previous study. We are inclined to conclude that they are homoplasmic after all. The observed heteroplasmy of *Mtr*BTN2 in Argentina may be due to double infection of mussels with *Mtr*BTN2 clones marked by different CR haplotypes or by somatic hybridization (cell fusion) between these clones. We believe that at least three different clones of *Mtr*BTN (one clone of *Mtr*BTN1 and two clones of *Mtr*BTN2) are parasitizing mussels in Magadan.

An unusually high *Mtr*BTN diversity in Magadan may be a matter of chance. Magadan is a port city, and infection could have been brought there by mussel-fouled ships from anywhere. On the other hand, Magadan is in the heart of the ancestral range of *M. trossulus* in the Northwest Pacific. *M. trossulus* populations in the Sea of Okhotsk are very abundant (Ushakov 1953; Khalaman et al. 2020) and presumably old. They might well turn out to be the natural reservoir of *M. trossulus*-derived transmissible cancers. Skazina et al. 2021 suggested that the Northwest Pacific may be a natural reservoir of *Mtr*BTN2. Now we can propose the same hypothesis for *Mtr*BTN1.

### Rapid test for BTN

A high labour-intensity of the methods used to diagnose *Mtr*BTN is the main reason why so little is known about the diversity and geographic distribution of this disease and about its prevalence in mussel populations. So far, only Hammel et al. (2021) have tried to address these issues comprehensively. They have identified *Mtr*BTN genotypes in a large sample of mussels (*M. edulis*, *M. galloprovincialis* and their hybrids) from Western Europe with the help of multilocus SNP genotyping using KASP (Kompetitive Allele Specific PCR). This method, based on competition of fluorescence of the two alleles, makes possible a quantitative assessment of the proportion of alternative alleles at a locus in a tissue sample. Chimeric genotypes of cancerous mussels are betrayed by an imbalanced proportion of the alleles, differing from that expected for homo- and heterozygotes.

This method is especially efficient for identification of *Mtr*BTN (which has the genotype of *M. trossulus*) in mussel species other than *M. trossulus* and also if many taxonomically informative markers are employed such as the SNP set used by Hammel et al. 2021. For KASP genotyping to be effective in the search for *Mtr*BTN in *M. trossulus* populations, more diagnostic markers for the cancer are needed.

In this study we used a rapid test of BTN based on parallel COI genotyping of the hemolymph and the muscle tissues. This method, though probably less sensitive than that employed by Hammel et al. (2021), is also less expensive. The test did not reveal known *Mtr*BTN alleles in the mussels with a rate of aneuploid cells less than 15% (Table 1). So, the suggested limit of its sensitivity is 15% of aneuploid cells in the hemolymph of a *Mtr*BTN-infected mussel.

The test demonstrated both heteroplasmy and its tissue-specific patterns in all mussels with *Mtr*BTN alleles except the most severely infected one (M2-22, 99% of aneuploid cells), which lacked the host alleles altogether. This means that not only known but also unknown cancerous alleles may potentially be detected using this test. We emphasize that a parallel analysis of the two tissues also makes it possible to distinguish between the cases of heteroplasmy related to chimerism due to BTN and the cases unrelated to it. In the former cases, the test would reveal differences between the foot and the hemolymph tissues, while in the latter cases, such as natural heteroplasmy or NUMTs co-amplification, it is unlikely that any such differences would be found.

Two mussels (M1-167 and M1-96) were a stumbling block in the interpretation of the test results. They had a high proportion of aneuploid cells in the hemolymph, no known cancerous alleles and no signs of genetic chimerism. Theoretically, these mussels could be infected by BTN lineages with mutations in the COI universal primers binding sites, which prevent its amplification. Alternatively, they may have conventional (non-transmissible) DN. Mussels with DN but without evidence of genetic chimerism have been repeatedly reported in *Mtr*BTN studies (Metzger et al. 2016; Burioli et al. 2019; Yonemitsu et al. 2019; Hammel et al. 2021). Hammel et al. 2021 made a special effort to prove that mussels can be affected with non-transmissible DN. Unfortunately, we did not study M1-167 and M1-96 for other markers and did not perform their molecular cloning. We do consider it likely that they had conventional DN, but this diagnosis is preliminary.

### Comparison between cancers

Different cancers of mussels from Magadan were quite similar in terms of ploidy (range 4.0-5.8n) and NCR (range of mean values 1.8-2.1). There were several genetic types of DN in our sample (supposedly non-transmissible neoplasia, *Mtr*BTN1 and two distinct clones of *Mtr*BTN2 marked by different COI haplotypes), and each type was represented by a few cases. The ranges of ploidy and NCR for these different DN types overlapped to a large extent. However, the ploidy of *Mtr*BTN2 in the Sea of Japan was somewhat lower (range 4.3n-4.9n) than in the Sea of Okhotsk (range 4.8n-5.8n). Interestingly, the two cases of presumably non-transmissible neoplasia were the most unusual in all the material; more interesting still, they were unusual for different reasons. In M1-167 the ploidy of the cancer cells was the lowest and strictly equal to four, which corresponds to tetraploidy. In M1-96, the cancer cells were noticeably smaller than in the other mussels (Table 1, 2, Supplementary Figure S2).

The relative uniformity of DN from the Sea of Okhotsk is especially noticeable when compared with the available data on DN in *M. edulis* from Western Europe, where *Mtr*BTN2 occurs, and in *M. trossulus* from the Northeastern Pacific, where *MtrBTN1* was registered. The survey of Burioli et al. (2019) showed that in Europe the ploidy of neoplastic cells varied in different cancerous mussels in the range of 8.36–18.8n; out of the two genetically confirmed cases of *Mtr*BTN, one had a ploidy of 8.4n and the other, with two peaks of aneuploid cells, 9.5n and 11.4n. They also observed that in different mussels the size of the nucleus of the neoplastic cells correlated positively with their ploidy. Hammel et al. (2021) compared genetically confirmed cases of *Mtr*BTN2 and conventional cancer from Europe (the transmissible and non-transmissible cancers were from different populations) on histological slides and recorded differences in the size of the nuclei, with the nuclei being about 40% larger in the transmissible cancers. In Northeastern Pacific (Washington and British Columbia), earlier studies reported tetraploid and pentaploid cancers (Elston et al. 1990; Moore et al. 1991), while a later study (Vassilenko and Baldwin 2014) reported a range of ploidy 1.4-5.5n of neoplastic cells between cancerous mussels and also within some mussels. As these studies preceded the discovery of BTN, the etiology of the disease is uncertain.

Thus, the ranges of ploidy values in mussels with neoplasia from Europe (*Mtr*BTN2-infected populations of *M. edulis*) and from Northeastern Pacific (*Mtr*BTN1-infected populations of *M. trossulus*) do not overlap, though in both regions the range is fairly broad. The range of ploidy values in mussels with neoplasia from Magadan (both lineages in *M. trossulus*, indistinguishable by ploidy) is narrow and almost overlaps with that of mussels in the Northeastern Pacific. A possible explanation of the ‘conservatism’ of cancers from the Sea of Okhotsk might be that the ploidy of *Mtr*BTN is controlled not only by its own genotype but also by the genotype of the host and its environment.

## Author Contributions

M.S., N.O. and P.S. designed the study. K.R. organized mussel sampling M.M., L.F. and N.O. provided cytological and histological analyses. M.S. and I.D. provided molecular genetic analyses. M.S. carried out the bioinformatics analysis. M.S. and P.S. drafted the manuscript. All authors read, approved, and contributed to the final manuscript.

## Acknowledgments

We would like to thank Natalia Lentsman for English language editing of the manuscript and Anna Romanovich, Alexey Masharskiy, Dmitriy Samulenkov, Dmitriy Zaitsev and Tatyiana Zemskova (St. Petersburg State University Research Park) for technical help.

