## Supplementary Figures and Tables for "High genetic diversity of bivalve transmissible neoplasia in the blue mussel *Mytilus trossulus* Gould from the subarctic Sea of Okhotsk"

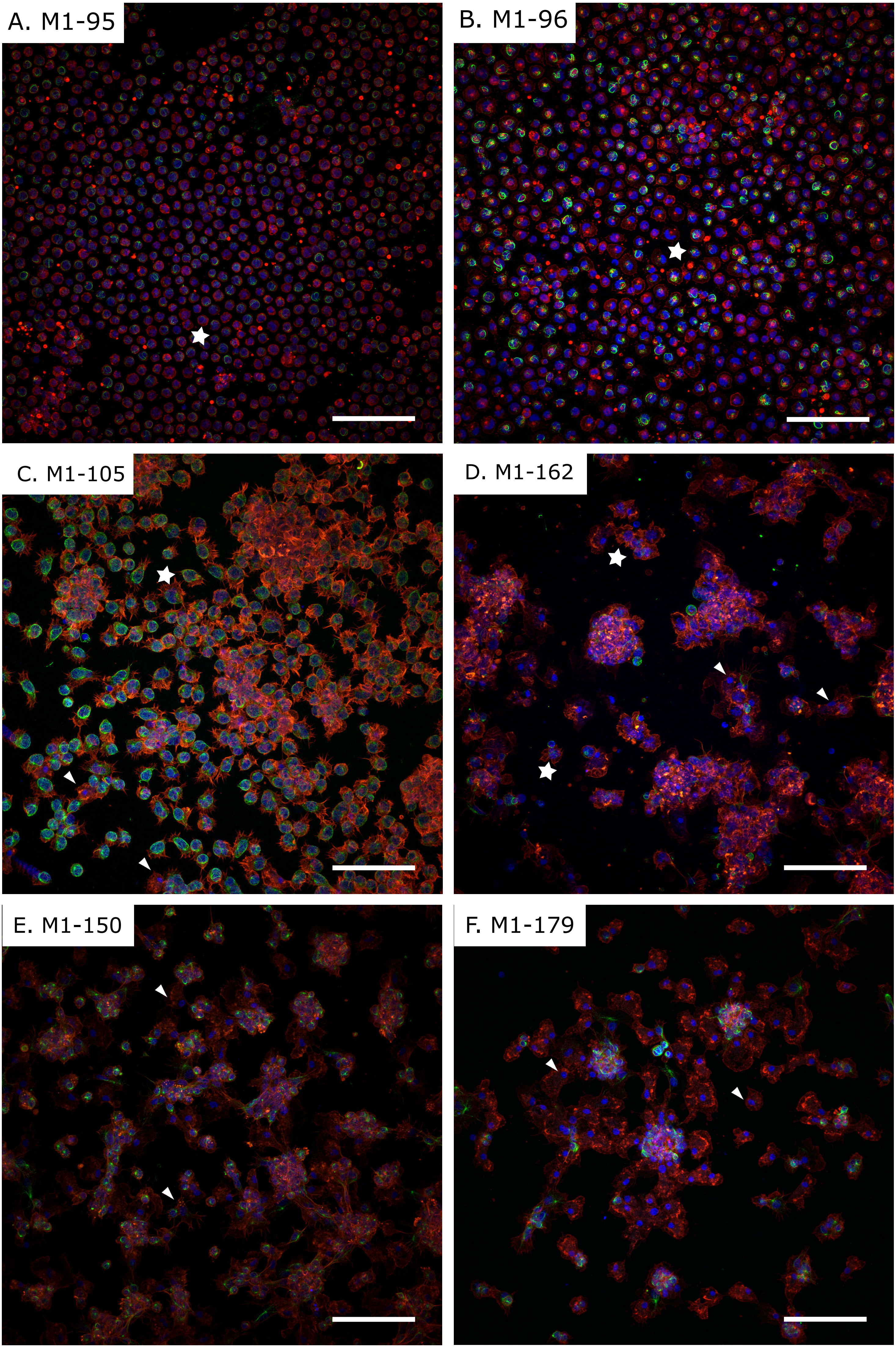

A. M1-95
B. M1-96
C. M1-105
D. M1-162
E. M1-150
F. M1-179

**Supplementary Figure S1**. Confocal microscopy images of hemocytes stained with DAPI (blue) TRITC-labelled phalloidin (red) and antibodies against tubulin (green) from DN-suggested mussels (**A-D**) and healthy control mussels (**E, F**). Arrowheads point to normally spread healthy hemocytes, while stars mark neoplastic hemocytes. Scale bar, 50 µm.

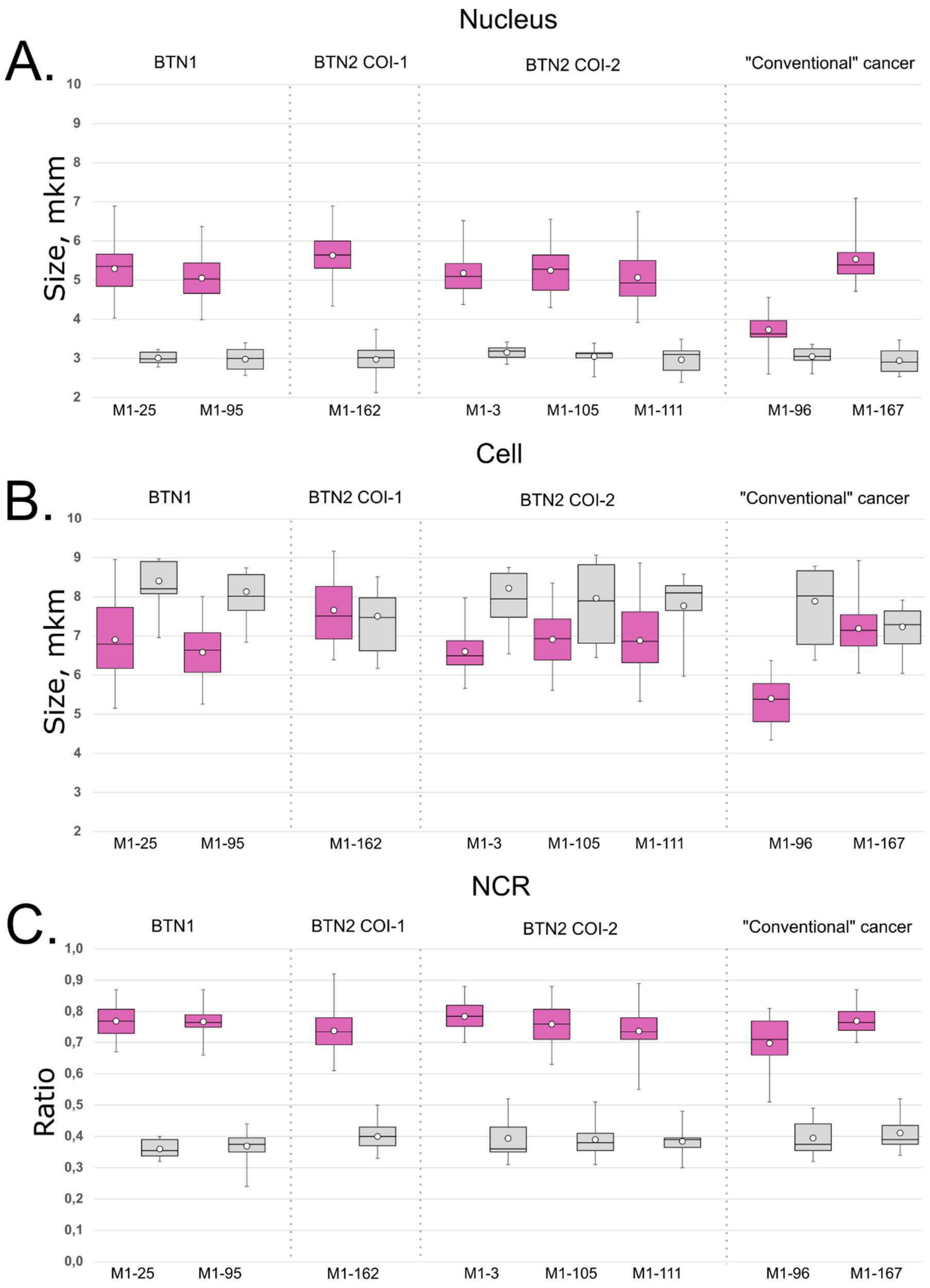


**Supplementary Figure S2.** Boxplots showing the range of variation for nucleus diameter (**A**), cell diameter (**B**), and NCR (**C**) for neoplastic hemocytes (violet boxes) and for healthy hemocytes (grey boxes). The boxplots represent the middle 50th percentile values. The horizontal line inside the boxplot indicates the median value for each category. The vertical lines indicate the upper and the lower ranges of values. The round marker inside the boxplot corresponds to the average value. Individuals are plotted along OX axes, size in µm or ratio (in C) is plotted along OX axes.

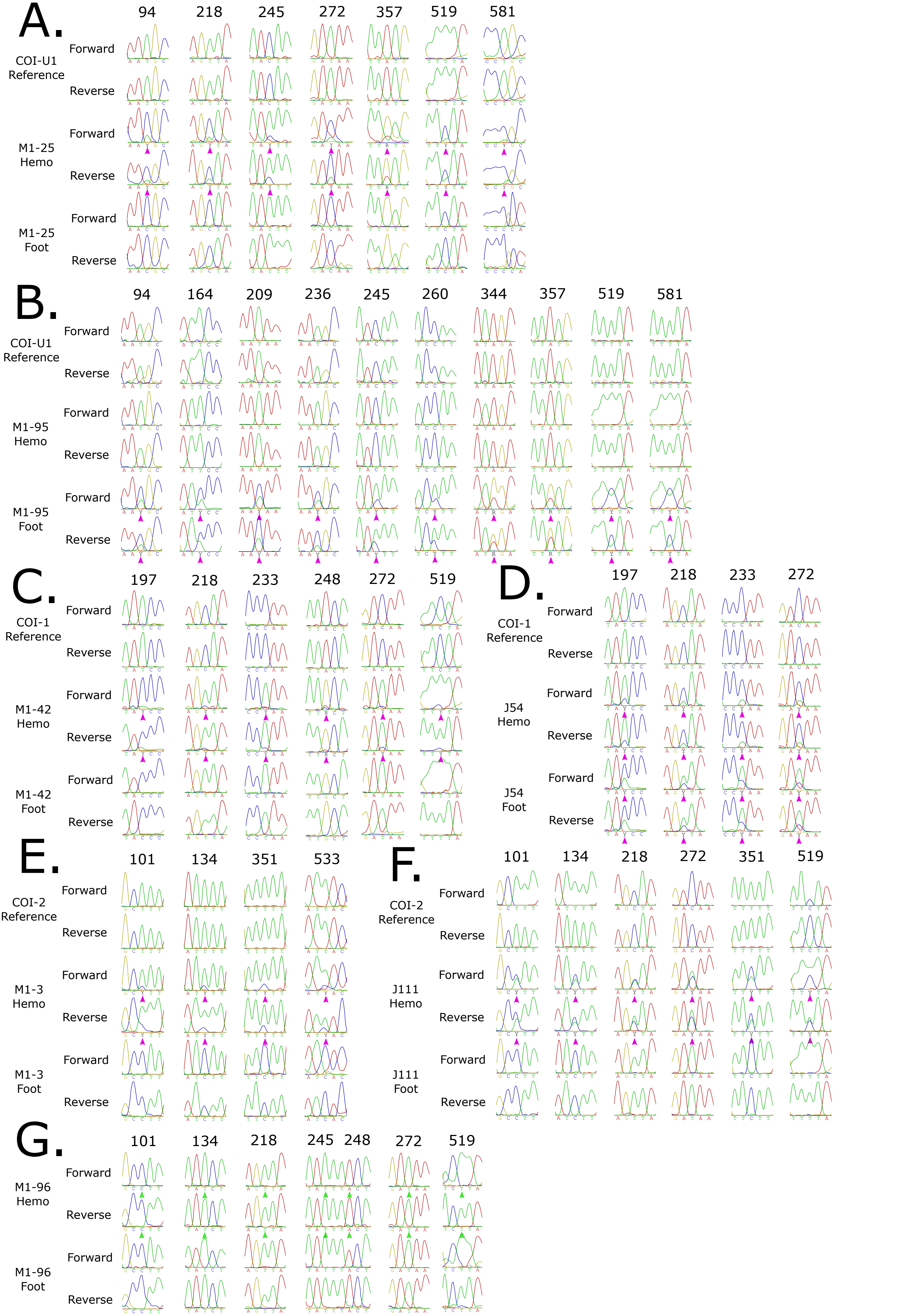

A.
94
218
245
272
357
519
581
COI-U1
Reference
Forward
Reverse
M1-25
Hemo
Forward
Reverse
M1-25
Foot
Forward
Reverse
B.
94
164
209
236
245
260
344
357
519
581
COI-U1
Reference
Forward
Reverse
M1-95
Hemo
Forward
Reverse
M1-95
Foot
Forward
Reverse
C.
197
218
233
248
272
519
COI-1
Reference
Forward
Reverse
M1-42
Hemo
Forward
Reverse
M1-42
Foot
Forward
Reverse
D.
197
218
233
272
COI-1
Reference
Forward
Reverse
J54
Hemo
Forward
Reverse
J54
Foot
Forward
Reverse
E.
101
134
351
533
COI-2
Reference
Forward
Reverse
M1-3
Hemo
Forward
Reverse
M1-3
Foot
Forward
Reverse
F.
101
134
218
272
351
519
COI-2
Reference
Forward
Reverse
J111
Hemo
Forward
Reverse
J111
Foot
Forward
Reverse
G.
101
134
218
245
248
272
519
M1-96
Hemo
Forward
Reverse
M1-96
Foot
Forward
Reverse

**Supplementary Figure S3.** Patterns of COI chromatograms observed in different tissues of Magadan and Gaydamak mussels: *Mtr*BTN-infected mussels M1-25 with 20% of neoplastic hemocytes (**A**), M1-95 with 99% (**B**), M1-42 with 15% (**C**), J54 with 44% (**D**), M1-3 with 45% (**E**), J111 with 26% (**F**), and M1-96, with 45% of aneuploid cells, allegedly having non-transmissible cancer (**G**). Numbers indicate nucleotide positions differing in host and cancer alleles according to the 630 bp alignment of COI used in this study. The first two lines (**A - F**) correspond to the reference cloned sequences of cancerous alleles, no reference is shown for M1-96 (**D**). The text block on the left provides individual ID, tissue of sequence origin, and the direction of sequencing. Violet triangles point to nucleotide positions with double peaks. Green triangles point to nucleotide positions without double peaks.

High-quality chromatograms with absent or low baseline noise make it possible to distinguish signals from minor additional sequences present in the PCR products of cancerous mussels. The following patterns observed in chromatograms from cancerous mussels are presented: peaks from the cancerous allele are minor in the hemolymph sequence and absent in the foot sequence (**A, C, E, F**); cancerous allele clearly dominates in hemolymph and is minor in foot tissues, as in highly infected individual M1-95 (**B**); double peaks are visible in both tissues (**D**). An individual without any double peaks is given as an example (**G**). Note that these patterns do not strongly depend on the level of neoplastic hemocytes estimated by flow cytometry. For example, J54 (**D**) and M1-3 (**E**) have almost identical levels of aneuploid cells in the hemolymph, but the cancerous allele is detected in the foot sequence in J54 and is absent in M1-3.

In chromatograms with double peaks, the genotypes of the host and the known cancerous lineages can be identified by the following signs. (1) Double peaks are observed in multiple positions indicating heteroplasmy. (2) The peaks from putatively cancer alleles are more pronounced in the hemolymph than in the foot tissues. (3) Nucleotide sequences of putatively cancer alleles correspond to known cancer alleles. (4) The above patterns can be recognized in both forward and reverse chromatograms.

Theoretically, in some positions, heteroplasmy could be imaginary. A minor signal could be residual from the major nucleotide of the same type right next to the position in question (e.g. minor peak from T in position 218 in chromatograms from the hemolymph of M1-25 could be residual from T in position 219). Because of the high quality of the chromatograms and the complete correspondence between the putative cancer alleles and the known cancer alleles, we do not think that the heteroplasmy at any position is false. However, since the quality of the chromatograms is high and since putative cancer alleles correspond completely to the known cancer alleles, we do not think that the heteroplasmy is false at any of the positions.

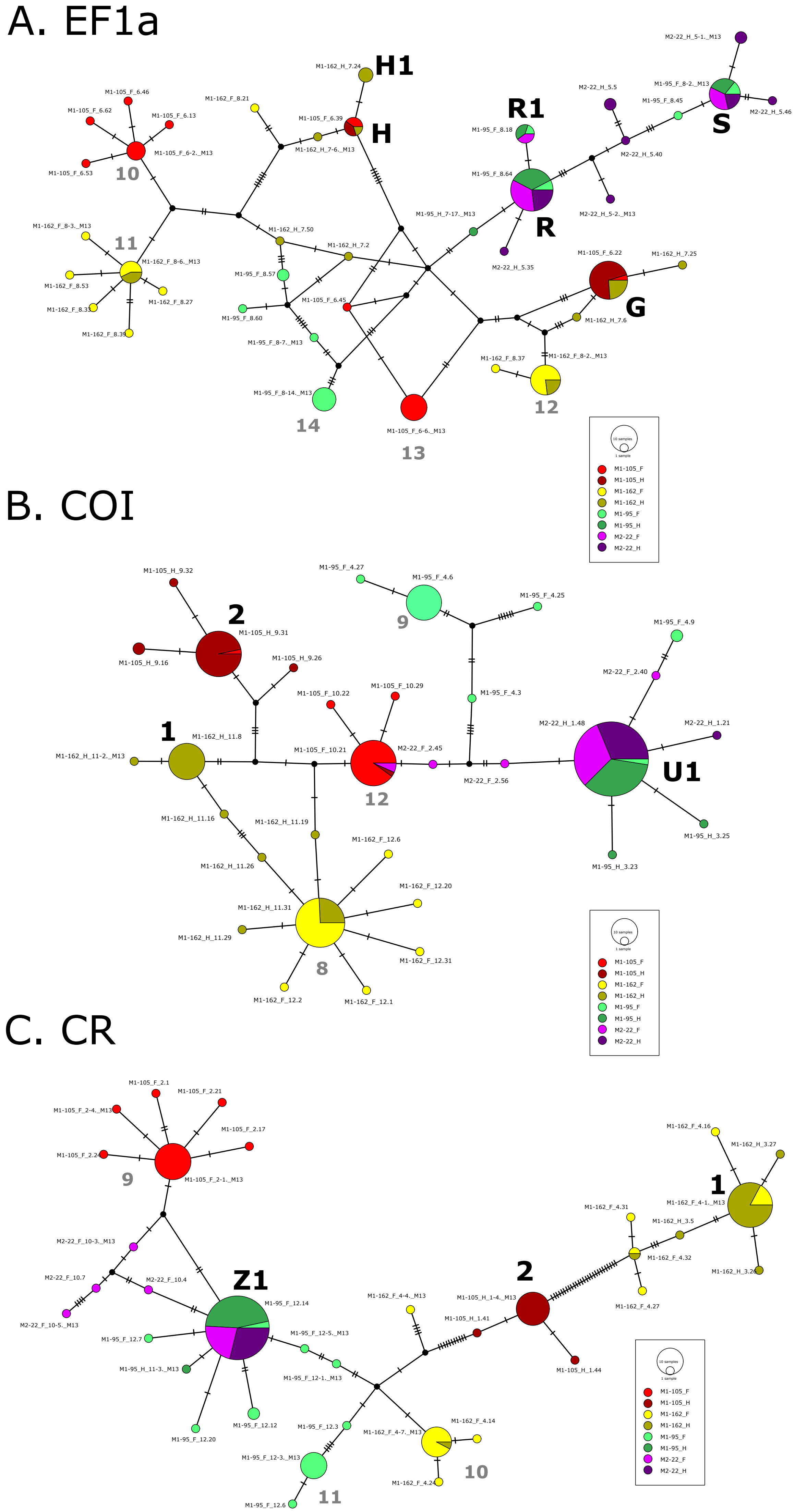

A. EF1a
H1
H
R1
R
S
G
10
11
12
13
14
M1-105_F_6.46
M1-105_F_6.62
M1-105_F_6.13
M1-105_F_6-2._M13
M1-105_F_6.53
M1-162_F_8.21
M1-105_F_6.39
M1-162_H_7.24
M1-162_H_7-6._M13
M1-162_F_8-3._M13
M1-162_F_8-6._M13
M1-162_F_8.53
M1-162_F_8.33
M1-162_F_8.27
M1-162_F_8.39
M1-162_H_7.50
M1-162_H_7.2
M1-95_F_8.57
M1-95_F_8.60
M1-105_F_6.45
M1-95_F_8-7._M13
M1-95_F_8-14._M13
M1-105_F_6-6._M13
M1-95_H_7-17._M13
M1-95_F_8.18
M1-95_F_8.64
M2-22_H_5.35
M2-22_H_5.5
M2-22_H_5.40
M2-22_H_5-2._M13
M2-22_H_5-1._M13
M1-95_F_8-2._M13
M1-95_F_8.45
M2-22_H_5.46
M1-105_F_6.22
M1-162_H_7.25
M1-162_H_7.6
M1-162_F_8.37
M1-162_F_8-2._M13
B. COI
2
1
9
12
U1
8
M1-105_H_9.32
M1-105_H_9.31
M1-105_H_9.16
M1-105_H_9.26
M1-95_F_4.27
M1-95_F_4.6
M1-95_F_4.25
M1-95_F_4.9
M1-95_F_4.3
M2-22_F_2.40
M1-105_F_10.22
M1-105_F_10.29
M1-105_F_10.21
M2-22_F_2.45
M2-22_F_2.56
M2-22_H_1.48
M2-22_H_1.21
M1-95_H_3.25
M1-95_H_3.23
M1-162_H_11.8
M1-162_H_11-2._M13
M1-162_H_11.16
M1-162_H_11.19
M1-162_H_11.26
M1-162_F_12.6
M1-162_F_12.20
M1-162_H_11.31
M1-162_H_11.29
M1-162_F_12.31
M1-162_F_12.2
M1-162_F_12.1
C. CR
9
Z1
2
1
10
11
M1-105_F_2.1
M1-105_F_2.21
M1-105_F_2-4._M13
M1-105_F_2.17
M1-105_F_2.24
M1-105_F_2-1._M13
M2-22_F_10-3._M13
M2-22_F_10.7
M2-22_F_10.4
M2-22_F_10-5._M13
M1-95_F_12.14
M1-95_F_12.7
M1-95_F_12-5._M13
M1-95_H_11-3._M13
M1-95_F_12-1._M13
M1-95_F_12.20
M1-95_F_12.12
M1-95_F_12.3
M1-95_F_12-3._M13
M1-95_F_12.6
M1-162_F_4-4._M13
M1-105_H_1-4._M13
M1-105_H_1.41
M1-105_H_1.44
M1-162_F_4-7._M13
M1-162_F_4.14
M1-162_F_4.24
M1-162_F_4.16
M1-162_H_3.27
M1-162_F_4-1._M13
M1-162_F_4.31
M1-162_H_3.5
M1-162_F_4.32
M1-162_F_4.27
M1-162_H_3.26
10 samples
1 sample
M1-105_F
M1-105_H
M1-162_F
M1-162_H
M1-95_F
M1-95_H
M2-22_F
M2-22_H

**Supplementary Figure S4.** TCS network representing all sequences obtained by molecular cloning of PCR fragments (**A**) EF1a, (**B**) COI and (**C**) CR. Sequences from individual mussels and from different tissues (hemolymph and foot) are colour-coded (see legend). Cancer haplotype IDs are given in black according to the nomenclature in Yonemitsu et al., 2019 and Skazina et al., 2021, host haplotype IDs are given in grey. Sequence IDs are given in small print, as in Supplementary Table S1. The variation in the poly-A region of CR was ignored as possibly arising from artefacts generated by polymerases in mononucleotide regions (Clarke et al, 2001).

Table S1. List of all sequences detected by molecular cloning of PCR products from different tissues of cancerous Magadan mussels

| Locus | Individual | Tissue | Clone | Haplotype |
|---|---|---|---|---|
| COI | M1-105 | Foot | 8 | COI-2 |
| COI | M1-105 | Foot | 1_M13 | COI-12 |
| COI | M1-105 | Foot | 2_M13 | COI-12 |
| COI | M1-105 | Foot | 4_M13 | COI-12 |
| COI | M1-105 | Foot | 6_M13 | COI-12 |
| COI | M1-105 | Foot | 1 | COI-12 |
| COI | M1-105 | Foot | 2 | COI-12 |
| COI | M1-105 | Foot | 3 | COI-12 |
| COI | M1-105 | Foot | 4 | COI-12 |
| COI | M1-105 | Foot | 5 | COI-12 |
| COI | M1-105 | Foot | 7 | COI-12 |
| COI | M1-105 | Foot | 10 | COI-12 |
| COI | M1-105 | Foot | 11 | COI-12 |
| COI | M1-105 | Foot | 13 | COI-12 |
| COI | M1-105 | Foot | 15 | COI-12 |
| COI | M1-105 | Foot | 13 | COI-12 |
| COI | M1-105 | Foot | 17 | COI-12 |
| COI | M1-105 | Foot | 18 | COI-12 |
| COI | M1-105 | Foot | 19 | COI-12 |
| COI | M1-105 | Foot | 27 | COI-12 |
| COI | M1-105 | Foot | 21 | COI-12 |
| COI | M1-105 | Foot | 23 | COI-12 |
| COI | M1-105 | Foot | 24 | COI-12 |
| COI | M1-105 | Foot | 26 | COI-12 |
| COI | M1-105 | Foot | 28 | COI-12 |
| COI | M1-105 | Foot | 30 | COI-12 |
| COI | M1-105 | Foot | 31 | COI-12 |
| COI | M1-105 | Foot | 32 | COI-12 |
| COI | M1-105 | Foot | 22 | MINOR |
| COI | M1-105 | Foot | 29 | MINOR |
| COI | M1-105 | Hemo | 5_M13 | COI-2 |
| COI | M1-105 | Hemo | 6_M13 | COI-2 |
| COI | M1-105 | Hemo | 7_M13 | COI-2 |
| COI | M1-105 | Hemo | 8_M13 | COI-2 |
| COI | M1-105 | Hemo | 1 | COI-2 |
| COI | M1-105 | Hemo | 2 | COI-2 |
| COI | M1-105 | Hemo | 3 | COI-2 |
| COI | M1-105 | Hemo | 4 | COI-2 |
| COI | M1-105 | Hemo | 5 | COI-2 |
| COI | M1-105 | Hemo | 6 | COI-2 |
| COI | M1-105 | Hemo | 7 | COI-2 |
| COI | M1-105 | Hemo | 9 | COI-2 |
| COI | M1-105 | Hemo | 10 | COI-2 |
| COI | M1-105 | Hemo | 11 | COI-2 |
| COI | M1-105 | Hemo | 13 | COI-2 |
| COI | M1-105 | Hemo | 14 | COI-2 |
| COI | M1-105 | Hemo | 15 | COI-2 |
| COI | M1-105 | Hemo | 17 | COI-2 |
| COI | M1-105 | Hemo | 18 | COI-2 |
| COI | M1-105 | Hemo | 19 | COI-2 |
| COI | M1-105 | Hemo | 20 | COI-2 |
| COI | M1-105 | Hemo | 21 | COI-2 |
| COI | M1-105 | Hemo | 22 | COI-2 |
| COI | M1-105 | Hemo | 23 | COI-2 |
| COI | M1-105 | Hemo | 24 | COI-2 |
| COI | M1-105 | Hemo | 25 | COI-2 |
| COI | M1-105 | Hemo | 28 | COI-2 |
| COI | M1-105 | Hemo | 29 | COI-2 |

| | | | | |
|---|---|---|---|---|
| COI | M1-105 | Hemo | 31 | COI-2 |
| COI | M1-105 | Hemo | 8 | COI-12 |
| COI | M1-105 | Hemo | 12 | MINOR |
| COI | M1-105 | Hemo | 16 | MINOR |
| COI | M1-105 | Hemo | 26 | MINOR |
| COI | M1-105 | Hemo | 32 | MINOR |
| COI | M1-162 | Foot | 3 | COI-8 |
| COI | M1-162 | Foot | 4 | COI-8 |
| COI | M1-162 | Foot | 5 | COI-8 |
| COI | M1-162 | Foot | 7 | COI-8 |
| COI | M1-162 | Foot | 8 | COI-8 |
| COI | M1-162 | Foot | 9 | COI-8 |
| COI | M1-162 | Foot | 10 | COI-8 |
| COI | M1-162 | Foot | 11 | COI-8 |
| COI | M1-162 | Foot | 12 | COI-8 |
| COI | M1-162 | Foot | 14 | COI-8 |
| COI | M1-162 | Foot | 15 | COI-8 |
| COI | M1-162 | Foot | 17 | COI-8 |
| COI | M1-162 | Foot | 18 | COI-8 |
| COI | M1-162 | Foot | 21 | COI-8 |
| COI | M1-162 | Foot | 24 | COI-8 |
| COI | M1-162 | Foot | 25 | COI-8 |
| COI | M1-162 | Foot | 26 | COI-8 |
| COI | M1-162 | Foot | 27 | COI-8 |
| COI | M1-162 | Foot | 28 | COI-8 |
| COI | M1-162 | Foot | 29 | COI-8 |
| COI | M1-162 | Foot | 30 | COI-8 |
| COI | M1-162 | Foot | 32 | COI-8 |
| COI | M1-162 | Foot | 1_M13 | COI-8 |
| COI | M1-162 | Foot | 2_M13 | COI-8 |
| COI | M1-162 | Foot | 3_M13 | COI-8 |
| COI | M1-162 | Foot | 4_M13 | COI-8 |
| COI | M1-162 | Foot | 1 | MINOR |
| COI | M1-162 | Foot | 2 | MINOR |
| COI | M1-162 | Foot | 6 | MINOR |
| COI | M1-162 | Foot | 20 | MINOR |
| COI | M1-162 | Foot | 31 | MINOR |
| COI | M1-162 | Hemo | 1_M13 | COI-1 |
| COI | M1-162 | Hemo | 3_M13 | COI-1 |
| COI | M1-162 | Hemo | 1 | COI-1 |
| COI | M1-162 | Hemo | 5 | COI-1 |
| COI | M1-162 | Hemo | 6 | COI-1 |
| COI | M1-162 | Hemo | 8 | COI-1 |
| COI | M1-162 | Hemo | 10 | COI-1 |
| COI | M1-162 | Hemo | 11 | COI-1 |
| COI | M1-162 | Hemo | 12 | COI-1 |
| COI | M1-162 | Hemo | 13 | COI-1 |
| COI | M1-162 | Hemo | 14 | COI-1 |
| COI | M1-162 | Hemo | 17 | COI-1 |
| COI | M1-162 | Hemo | 18 | COI-1 |
| COI | M1-162 | Hemo | 21 | COI-1 |
| COI | M1-162 | Hemo | 22 | COI-1 |
| COI | M1-162 | Hemo | 24 | COI-1 |
| COI | M1-162 | Hemo | 27 | COI-1 |
| COI | M1-162 | Hemo | 30 | COI-1 |
| COI | M1-162 | Hemo | 32 | COI-1 |
| COI | M1-162 | Hemo | 4_M13 | COI-8 |
| COI | M1-162 | Hemo | 2 | COI-8 |
| COI | M1-162 | Hemo | 3 | COI-8 |
| COI | M1-162 | Hemo | 15 | COI-8 |
| COI | M1-162 | Hemo | 20 | COI-8 |
| COI | M1-162 | Hemo | 23 | COI-8 |
| COI | M1-162 | Hemo | 25 | COI-8 |
| COI | M1-162 | Hemo | 28 | COI-8 |
| COI | M1-162 | Hemo | 31 | COI-8 |

| | | | | |
|---|---|---|---|---|
| COI | M1-162 | Hemo | 2_M13 | MINOR |
| COI | M1-162 | Hemo | 16 | MINOR |
| COI | M1-162 | Hemo | 19 | MINOR |
| COI | M1-162 | Hemo | 26 | MINOR |
| COI | M1-162 | Hemo | 29 | MINOR |
| COI | M1-95 | Foot | 2 | COI-9 |
| COI | M1-95 | Foot | 6 | COI-9 |
| COI | M1-95 | Foot | 10 | COI-9 |
| COI | M1-95 | Foot | 12 | COI-9 |
| COI | M1-95 | Foot | 13 | COI-9 |
| COI | M1-95 | Foot | 15 | COI-9 |
| COI | M1-95 | Foot | 16 | COI-9 |
| COI | M1-95 | Foot | 19 | COI-9 |
| COI | M1-95 | Foot | 21 | COI-9 |
| COI | M1-95 | Foot | 23 | COI-9 |
| COI | M1-95 | Foot | 24 | COI-9 |
| COI | M1-95 | Foot | 26 | COI-9 |
| COI | M1-95 | Foot | 28 | COI-9 |
| COI | M1-95 | Foot | 30 | COI-9 |
| COI | M1-95 | Foot | 31 | COI-9 |
| COI | M1-95 | Foot | 1_M13 | COI-9 |
| COI | M1-95 | Foot | 4_M13 | COI-9 |
| COI | M1-95 | Foot | 5_M13 | COI-9 |
| COI | M1-95 | Foot | 3 | MINOR |
| COI | M1-95 | Foot | 9 | MINOR |
| COI | M1-95 | Foot | 11 | MINOR |
| COI | M1-95 | Foot | 25 | MINOR |
| COI | M1-95 | Foot | 27 | MINOR |
| COI | M1-95 | Foot | 4 | U1 |
| COI | M1-95 | Foot | 6_M13 | U1 |
| COI | M1-95 | Hemo | 23 | MINOR |
| COI | M1-95 | Hemo | 25 | MINOR |
| COI | M1-95 | Hemo | 1_M13 | U1 |
| COI | M1-95 | Hemo | 2_M13 | U1 |
| COI | M1-95 | Hemo | 3_M13 | U1 |
| COI | M1-95 | Hemo | 4_M13 | U1 |
| COI | M1-95 | Hemo | 1 | U1 |
| COI | M1-95 | Hemo | 2 | U1 |
| COI | M1-95 | Hemo | 3 | U1 |
| COI | M1-95 | Hemo | 5 | U1 |
| COI | M1-95 | Hemo | 6 | U1 |
| COI | M1-95 | Hemo | 8 | U1 |
| COI | M1-95 | Hemo | 9 | U1 |
| COI | M1-95 | Hemo | 11 | U1 |
| COI | M1-95 | Hemo | 12 | U1 |
| COI | M1-95 | Hemo | 13 | U1 |
| COI | M1-95 | Hemo | 14 | U1 |
| COI | M1-95 | Hemo | 15 | U1 |
| COI | M1-95 | Hemo | 16 | U1 |
| COI | M1-95 | Hemo | 17 | U1 |
| COI | M1-95 | Hemo | 18 | U1 |
| COI | M1-95 | Hemo | 19 | U1 |
| COI | M1-95 | Hemo | 20 | U1 |
| COI | M1-95 | Hemo | 22 | U1 |
| COI | M1-95 | Hemo | 24 | U1 |
| COI | M1-95 | Hemo | 26 | U1 |
| COI | M1-95 | Hemo | 27 | U1 |
| COI | M1-95 | Hemo | 29 | U1 |
| COI | M1-95 | Hemo | 30 | U1 |
| COI | M1-95 | Hemo | 32 | U1 |
| COI | M2-22 | Foot | 2_M13 | COI-12 |
| COI | M2-22 | Foot | 5_M13 | COI-12 |
| COI | M2-22 | Foot | 40 | MINOR |
| COI | M2-22 | Foot | 45 | MINOR |
| COI | M2-22 | Foot | 56 | MINOR |

| | | | | |
|---|---|---|---|---|
| COI | M2-22 | Foot | 3_M13 | COI-U1 |
| COI | M2-22 | Foot | 4_M13 | COI-U1 |
| COI | M2-22 | Foot | 33 | COI-U1 |
| COI | M2-22 | Foot | 35 | COI-U1 |
| COI | M2-22 | Foot | 36 | COI-U1 |
| COI | M2-22 | Foot | 38 | COI-U1 |
| COI | M2-22 | Foot | 42 | COI-U1 |
| COI | M2-22 | Foot | 43 | COI-U1 |
| COI | M2-22 | Foot | 44 | COI-U1 |
| COI | M2-22 | Foot | 47 | COI-U1 |
| COI | M2-22 | Foot | 48 | COI-U1 |
| COI | M2-22 | Foot | 49 | COI-U1 |
| COI | M2-22 | Foot | 50 | COI-U1 |
| COI | M2-22 | Foot | 51 | COI-U1 |
| COI | M2-22 | Foot | 52 | COI-U1 |
| COI | M2-22 | Foot | 53 | COI-U1 |
| COI | M2-22 | Foot | 54 | COI-U1 |
| COI | M2-22 | Foot | 55 | COI-U1 |
| COI | M2-22 | Foot | 57 | COI-U1 |
| COI | M2-22 | Foot | 58 | COI-U1 |
| COI | M2-22 | Foot | 59 | COI-U1 |
| COI | M2-22 | Foot | 61 | COI-U1 |
| COI | M2-22 | Foot | 62 | COI-U1 |
| COI | M2-22 | Foot | 63 | COI-U1 |
| COI | M2-22 | Foot | 64 | COI-U1 |
| COI | M2-22 | Hemo | 21 | MINOR |
| COI | M2-22 | Hemo | 1_M13 | COI-U1 |
| COI | M2-22 | Hemo | 2_M13 | COI-U1 |
| COI | M2-22 | Hemo | 3_M13 | COI-U1 |
| COI | M2-22 | Hemo | 4_M13 | COI-U1 |
| COI | M2-22 | Hemo | 3 | COI-U1 |
| COI | M2-22 | Hemo | 4 | COI-U1 |
| COI | M2-22 | Hemo | 9 | COI-U1 |
| COI | M2-22 | Hemo | 10 | COI-U1 |
| COI | M2-22 | Hemo | 12 | COI-U1 |
| COI | M2-22 | Hemo | 13 | COI-U1 |
| COI | M2-22 | Hemo | 15 | COI-U1 |
| COI | M2-22 | Hemo | 19 | COI-U1 |
| COI | M2-22 | Hemo | 28 | COI-U1 |
| COI | M2-22 | Hemo | 33 | COI-U1 |
| COI | M2-22 | Hemo | 35 | COI-U1 |
| COI | M2-22 | Hemo | 36 | COI-U1 |
| COI | M2-22 | Hemo | 37 | COI-U1 |
| COI | M2-22 | Hemo | 39 | COI-U1 |
| COI | M2-22 | Hemo | 40 | COI-U1 |
| COI | M2-22 | Hemo | 41 | COI-U1 |
| COI | M2-22 | Hemo | 42 | COI-U1 |
| COI | M2-22 | Hemo | 43 | COI-U1 |
| COI | M2-22 | Hemo | 45 | COI-U1 |
| COI | M2-22 | Hemo | 46 | COI-U1 |
| COI | M2-22 | Hemo | 48 | COI-U1 |
| CR | M1-105 | Foot | 2 | CR-9 |
| CR | M1-105 | Foot | 4 | CR-9 |
| CR | M1-105 | Foot | 6 | CR-9 |
| CR | M1-105 | Foot | 8 | CR-9 |
| CR | M1-105 | Foot | 9 | CR-9 |
| CR | M1-105 | Foot | 11 | CR-9 |
| CR | M1-105 | Foot | 12 | CR-9 |
| CR | M1-105 | Foot | 13 | CR-9 |
| CR | M1-105 | Foot | 14 | CR-9 |
| CR | M1-105 | Foot | 16 | CR-9 |
| CR | M1-105 | Foot | 19 | CR-9 |
| CR | M1-105 | Foot | 20 | CR-9 |
| CR | M1-105 | Foot | 25 | CR-9 |
| CR | M1-105 | Foot | 27 | CR-9 |

| | | | | |
|---|---|---|---|---|
| CR | M1-105 | Foot | 29 | CR-9 |
| CR | M1-105 | Foot | 31 | CR-9 |
| CR | M1-105 | Foot | 1_M13 | CR-9 |
| CR | M1-105 | Foot | 2_M13 | CR-9 |
| CR | M1-105 | Foot | 7_M13 | CR-9 |
| CR | M1-105 | Foot | 1 | MINOR |
| CR | M1-105 | Foot | 17 | MINOR |
| CR | M1-105 | Foot | 21 | MINOR |
| CR | M1-105 | Foot | 24 | MINOR |
| CR | M1-105 | Foot | 4_M13 | MINOR |
| CR | M1-105 | Hemo | 4_M13 | CR-2 |
| CR | M1-105 | Hemo | 5_M13 | CR-2 |
| CR | M1-105 | Hemo | 6_M13 | CR-2 |
| CR | M1-105 | Hemo | 8_M13 | CR-2 |
| CR | M1-105 | Hemo | 3 | CR-2 |
| CR | M1-105 | Hemo | 5 | CR-2 |
| CR | M1-105 | Hemo | 6 | CR-2 |
| CR | M1-105 | Hemo | 7 | CR-2 |
| CR | M1-105 | Hemo | 10 | CR-2 |
| CR | M1-105 | Hemo | 13 | CR-2 |
| CR | M1-105 | Hemo | 17 | CR-2 |
| CR | M1-105 | Hemo | 21 | CR-2 |
| CR | M1-105 | Hemo | 29 | CR-2 |
| CR | M1-105 | Hemo | 35 | CR-2 |
| CR | M1-105 | Hemo | 36 | CR-2 |
| CR | M1-105 | Hemo | 39 | CR-2 |
| CR | M1-105 | Hemo | 41 | MINOR |
| CR | M1-105 | Hemo | 44 | MINOR |
| CR | M1-162 | Foot | 2 | CR-1 |
| CR | M1-162 | Foot | 7 | CR-1 |
| CR | M1-162 | Foot | 29 | CR-1 |
| CR | M1-162 | Foot | 1_M13 | CR-1 |
| CR | M1-162 | Foot | 3_M13 | CR-1 |
| CR | M1-162 | Foot | 1 | CR-10 |
| CR | M1-162 | Foot | 3 | CR-10 |
| CR | M1-162 | Foot | 4 | CR-10 |
| CR | M1-162 | Foot | 5 | CR-10 |
| CR | M1-162 | Foot | 9 | CR-10 |
| CR | M1-162 | Foot | 11 | CR-10 |
| CR | M1-162 | Foot | 12 | CR-10 |
| CR | M1-162 | Foot | 18 | CR-10 |
| CR | M1-162 | Foot | 20 | CR-10 |
| CR | M1-162 | Foot | 23 | CR-10 |
| CR | M1-162 | Foot | 26 | CR-10 |
| CR | M1-162 | Foot | 30 | CR-10 |
| CR | M1-162 | Foot | 7_M13 | CR-10 |
| CR | M1-162 | Foot | 14 | MINOR |
| CR | M1-162 | Foot | 16 | MINOR |
| CR | M1-162 | Foot | 24 | MINOR |
| CR | M1-162 | Foot | 27 | MINOR |
| CR | M1-162 | Foot | 31 | MINOR |
| CR | M1-162 | Foot | 32 | MINOR |
| CR | M1-162 | Foot | 4_M13 | MINOR |
| CR | M1-162 | Hemo | 1_M13 | CR-1 |
| CR | M1-162 | Hemo | 3_M13 | CR-1 |
| CR | M1-162 | Hemo | 4_M13 | CR-1 |
| CR | M1-162 | Hemo | 1 | CR-1 |
| CR | M1-162 | Hemo | 3 | CR-1 |
| CR | M1-162 | Hemo | 4 | CR-1 |
| CR | M1-162 | Hemo | 6 | CR-1 |
| CR | M1-162 | Hemo | 8 | CR-1 |
| CR | M1-162 | Hemo | 9 | CR-1 |
| CR | M1-162 | Hemo | 11 | CR-1 |
| CR | M1-162 | Hemo | 13 | CR-1 |
| CR | M1-162 | Hemo | 14 | CR-1 |

| | | | | |
|---|---|---|---|---|
| CR | M1-162 | Hemo | 15 | CR-1 |
| CR | M1-162 | Hemo | 16 | CR-1 |
| CR | M1-162 | Hemo | 17 | CR-1 |
| CR | M1-162 | Hemo | 19 | CR-1 |
| CR | M1-162 | Hemo | 21 | CR-1 |
| CR | M1-162 | Hemo | 22 | CR-1 |
| CR | M1-162 | Hemo | 23 | CR-1 |
| CR | M1-162 | Hemo | 24 | CR-1 |
| CR | M1-162 | Hemo | 25 | CR-1 |
| CR | M1-162 | Hemo | 28 | CR-1 |
| CR | M1-162 | Hemo | 29 | CR-1 |
| CR | M1-162 | Hemo | 30 | CR-1 |
| CR | M1-162 | Hemo | 31 | CR-1 |
| CR | M1-162 | Hemo | 5 | MINOR |
| CR | M1-162 | Hemo | 7 | MINOR |
| CR | M1-162 | Hemo | 26 | MINOR |
| CR | M1-162 | Hemo | 27 | MINOR |
| CR | M1-95 | Foot | 1 | CR-11 |
| CR | M1-95 | Foot | 8 | CR-11 |
| CR | M1-95 | Foot | 9 | CR-11 |
| CR | M1-95 | Foot | 10 | CR-11 |
| CR | M1-95 | Foot | 11 | CR-11 |
| CR | M1-95 | Foot | 13 | CR-11 |
| CR | M1-95 | Foot | 16 | CR-11 |
| CR | M1-95 | Foot | 19 | CR-11 |
| CR | M1-95 | Foot | 3_M13 | CR-11 |
| CR | M1-95 | Foot | 4_M13 | CR-11 |
| CR | M1-95 | Foot | 3 | MINOR |
| CR | M1-95 | Foot | 6 | MINOR |
| CR | M1-95 | Foot | 7 | MINOR |
| CR | M1-95 | Foot | 12 | MINOR |
| CR | M1-95 | Foot | 18 | MINOR |
| CR | M1-95 | Foot | 20 | MINOR |
| CR | M1-95 | Foot | 1_M13 | MINOR |
| CR | M1-95 | Foot | 5_M13 | MINOR |
| CR | M1-95 | Foot | 14 | CR-Z1 |
| CR | M1-95 | Foot | 22 | CR-Z1 |
| CR | M1-95 | Hemo | 3_M13 | MINOR |
| CR | M1-95 | Hemo | 18 | CR-Z1 |
| CR | M1-95 | Hemo | 22 | CR-Z1 |
| CR | M1-95 | Hemo | 11 | CR-Z1 |
| CR | M1-95 | Hemo | 9 | CR-Z1 |
| CR | M1-95 | Hemo | 12 | CR-Z1 |
| CR | M1-95 | Hemo | 5 | CR-Z1 |
| CR | M1-95 | Hemo | 8 | CR-Z1 |
| CR | M1-95 | Hemo | 14 | CR-Z1 |
| CR | M1-95 | Hemo | 7 | CR-Z1 |
| CR | M1-95 | Hemo | 10 | CR-Z1 |
| CR | M1-95 | Hemo | 6 | CR-Z1 |
| CR | M1-95 | Hemo | 24 | CR-Z1 |
| CR | M1-95 | Hemo | 1 | CR-Z1 |
| CR | M1-95 | Hemo | 15 | CR-Z1 |
| CR | M1-95 | Hemo | 23 | CR-Z1 |
| CR | M1-95 | Hemo | 3 | CR-Z1 |
| CR | M1-95 | Hemo | 16 | CR-Z1 |
| CR | M1-95 | Hemo | 19 | CR-Z1 |
| CR | M1-95 | Hemo | 20 | CR-Z1 |
| CR | M1-95 | Hemo | 21 | CR-Z1 |
| CR | M1-95 | Hemo | 13 | CR-Z1 |
| CR | M1-95 | Hemo | 17 | CR-Z1 |
| CR | M1-95 | Hemo | 4 | CR-Z1 |
| CR | M1-95 | Hemo | 2 | CR-Z1 |
| CR | M1-95 | Hemo | 1_M13 | CR-Z1 |
| CR | M1-95 | Hemo | 2_M13 | CR-Z1 |
| CR | M1-95 | Hemo | 6_M13 | CR-Z1 |

| | | | | |
|---|---|---|---|---|
| CR | M2-22 | Foot | 7 | MINOR |
| CR | M2-22 | Foot | 4 | MINOR |
| CR | M2-22 | Foot | 3_M13 | MINOR |
| CR | M2-22 | Foot | 5 | MINOR |
| CR | M2-22 | Foot | 6 | CR-Z1 |
| CR | M2-22 | Foot | 8 | CR-Z1 |
| CR | M2-22 | Foot | 2 | CR-Z1 |
| CR | M2-22 | Foot | 10 | CR-Z1 |
| CR | M2-22 | Foot | 1 | CR-Z1 |
| CR | M2-22 | Foot | 4_M13 | CR-Z1 |
| CR | M2-22 | Foot | 12 | CR-Z1 |
| CR | M2-22 | Foot | 21 | CR-Z1 |
| CR | M2-22 | Foot | 13 | CR-Z1 |
| CR | M2-22 | Foot | 1_M13 | CR-Z1 |
| CR | M2-22 | Foot | 11 | CR-Z1 |
| CR | M2-22 | Foot | 3 | CR-Z1 |
| CR | M2-22 | Hemo | 1_M13 | CR-Z1 |
| CR | M2-22 | Hemo | 5_M13 | CR-Z1 |
| CR | M2-22 | Hemo | 3_M13 | CR-Z1 |
| CR | M2-22 | Hemo | 2_M13 | CR-Z1 |
| CR | M2-22 | Hemo | 22 | CR-Z1 |
| CR | M2-22 | Hemo | 14 | CR-Z1 |
| CR | M2-22 | Hemo | 15 | CR-Z1 |
| CR | M2-22 | Hemo | 20 | CR-Z1 |
| CR | M2-22 | Hemo | 18 | CR-Z1 |
| CR | M2-22 | Hemo | 17 | CR-Z1 |
| CR | M2-22 | Hemo | 23 | CR-Z1 |
| CR | M2-22 | Hemo | 7 | CR-Z1 |
| CR | M2-22 | Hemo | 9 | CR-Z1 |
| CR | M2-22 | Hemo | 16 | CR-Z1 |
| CR | M2-22 | Hemo | 1 | CR-Z1 |
| CR | M2-22 | Hemo | 4 | CR-Z1 |
| CR | M2-22 | Hemo | 10 | CR-Z1 |
| EF1a | M1-105 | Foot | 22 | EF1a-G |
| EF1a | M1-105 | Foot | 39 | EF1a-H |
| EF1a | M1-105 | Foot | 49 | EF1a-H |
| EF1a | M1-105 | Foot | 24 | EF1a-10 |
| EF1a | M1-105 | Foot | 50 | EF1a-10 |
| EF1a | M1-105 | Foot | 2_M13 | EF1a-10 |
| EF1a | M1-105 | Foot | 4_M13 | EF1a-10 |
| EF1a | M1-105 | Foot | 64 | EF1a-10 |
| EF1a | M1-105 | Foot | 4 | EF1a-13 |
| EF1a | M1-105 | Foot | 7 | EF1a-13 |
| EF1a | M1-105 | Foot | 18 | EF1a-13 |
| EF1a | M1-105 | Foot | 33 | EF1a-13 |
| EF1a | M1-105 | Foot | 35 | EF1a-13 |
| EF1a | M1-105 | Foot | 40 | EF1a-13 |
| EF1a | M1-105 | Foot | 41 | EF1a-13 |
| EF1a | M1-105 | Foot | 47 | EF1a-13 |
| EF1a | M1-105 | Foot | 6_M13 | EF1a-13 |
| EF1a | M1-105 | Foot | 7_M13 | EF1a-13 |
| EF1a | M1-105 | Foot | 13 | MINOR |
| EF1a | M1-105 | Foot | 45 | MINOR |
| EF1a | M1-105 | Foot | 46 | MINOR |
| EF1a | M1-105 | Foot | 53 | MINOR |
| EF1a | M1-105 | Foot | 62 | MINOR |
| EF1a | M1-105 | Hemo | 2_M13 | EF1a-G |
| EF1a | M1-105 | Hemo | 7_M13 | EF1a-G |
| EF1a | M1-105 | Hemo | 4_M13 | EF1a-G |
| EF1a | M1-105 | Hemo | 16 | EF1a-G |
| EF1a | M1-105 | Hemo | 49 | EF1a-G |
| EF1a | M1-105 | Hemo | 29 | EF1a-G |
| EF1a | M1-105 | Hemo | 32 | EF1a-G |
| EF1a | M1-105 | Hemo | 12 | EF1a-G |
| EF1a | M1-105 | Hemo | 52 | EF1a-G |

| | | | | |
|---|---|---|---|---|
| EF1a | M1-105 | Hemo | 11 | EF1a-G |
| EF1a | M1-105 | Hemo | 30 | EF1a-G |
| EF1a | M1-105 | Hemo | 59 | EF1a-G |
| EF1a | M1-105 | Hemo | 40 | EF1a-G |
| EF1a | M1-105 | Hemo | 50 | EF1a-G |
| EF1a | M1-105 | Hemo | 19 | EF1a-G |
| EF1a | M1-105 | Hemo | 3_M13 | EF1a-H |
| EF1a | M1-105 | Hemo | 17 | EF1a-H |
| EF1a | M1-162 | Foot | 32 | EF1a-11 |
| EF1a | M1-162 | Foot | 38 | EF1a-11 |
| EF1a | M1-162 | Foot | 6_M13 | EF1a-11 |
| EF1a | M1-162 | Foot | 8_M13 | EF1a-11 |
| EF1a | M1-162 | Foot | 18 | EF1a-12 |
| EF1a | M1-162 | Foot | 23 | EF1a-12 |
| EF1a | M1-162 | Foot | 26 | EF1a-12 |
| EF1a | M1-162 | Foot | 28 | EF1a-12 |
| EF1a | M1-162 | Foot | 41 | EF1a-12 |
| EF1a | M1-162 | Foot | 45 | EF1a-12 |
| EF1a | M1-162 | Foot | 46 | EF1a-12 |
| EF1a | M1-162 | Foot | 48 | EF1a-12 |
| EF1a | M1-162 | Foot | 50 | EF1a-12 |
| EF1a | M1-162 | Foot | 2_M13 | EF1a-12 |
| EF1a | M1-162 | Foot | 21 | MINOR |
| EF1a | M1-162 | Foot | 27 | MINOR |
| EF1a | M1-162 | Foot | 33 | MINOR |
| EF1a | M1-162 | Foot | 37 | MINOR |
| EF1a | M1-162 | Foot | 39 | MINOR |
| EF1a | M1-162 | Foot | 53 | MINOR |
| EF1a | M1-162 | Foot | 3_M13 | MINOR |
| EF1a | M1-162 | Hemo | 5 | EF1a-G |
| EF1a | M1-162 | Hemo | 51 | EF1a-G |
| EF1a | M1-162 | Hemo | 60 | EF1a-G |
| EF1a | M1-162 | Hemo | 67 | EF1a-G |
| EF1a | M1-162 | Hemo | 4_M13 | EF1a-G |
| EF1a | M1-162 | Hemo | 66 | EF1a-H |
| EF1a | M1-162 | Hemo | 9 | EF1a-H1 |
| EF1a | M1-162 | Hemo | 24 | EF1a-H1 |
| EF1a | M1-162 | Hemo | 59 | EF1a-H1 |
| EF1a | M1-162 | Hemo | 11 | EF1a-11 |
| EF1a | M1-162 | Hemo | 17 | EF1a-11 |
| EF1a | M1-162 | Hemo | 18 | EF1a-11 |
| EF1a | M1-162 | Hemo | 1_M13 | EF1a-12 |
| EF1a | M1-162 | Hemo | 3_M13 | EF1a-12 |
| EF1a | M1-162 | Hemo | 5_M13 | EF1a-12 |
| EF1a | M1-162 | Hemo | 2 | MINOR |
| EF1a | M1-162 | Hemo | 6 | MINOR |
| EF1a | M1-162 | Hemo | 25 | MINOR |
| EF1a | M1-162 | Hemo | 50 | MINOR |
| EF1a | M1-162 | Hemo | 6_M13 | MINOR |
| EF1a | M1-95 | Foot | 14_M13 | EF1a-14 |
| EF1a | M1-95 | Foot | 21 | EF1a-14 |
| EF1a | M1-95 | Foot | 30 | EF1a-14 |
| EF1a | M1-95 | Foot | 51 | EF1a-14 |
| EF1a | M1-95 | Foot | 55 | EF1a-14 |
| EF1a | M1-95 | Foot | 59 | EF1a-14 |
| EF1a | M1-95 | Foot | 67 | EF1a-14 |
| EF1a | M1-95 | Foot | 68 | EF1a-14 |
| EF1a | M1-95 | Foot | 57 | MINOR |
| EF1a | M1-95 | Foot | 70 | MINOR |
| EF1a | M1-95 | Foot | 60 | MINOR |
| EF1a | M1-95 | Foot | 45 | MINOR |
| EF1a | M1-95 | Foot | 7_M13 | MINOR |
| EF1a | M1-95 | Foot | 64 | EF1a-R |
| EF1a | M1-95 | Foot | 69 | EF1a-R |
| EF1a | M1-95 | Foot | 18 | EF1a-R1 |

| | | | | |
|---|---|---|---|---|
| EF1a | M1-95 | Foot | 2_M13 | EF1a-S |
| EF1a | M1-95 | Foot | 5_M13 | EF1a-S |
| EF1a | M1-95 | Hemo | 17_M13 | MINOR |
| EF1a | M1-95 | Hemo | 1_M13 | EF1a-R |
| EF1a | M1-95 | Hemo | 3_M13 | EF1a-R |
| EF1a | M1-95 | Hemo | 6_M13 | EF1a-R |
| EF1a | M1-95 | Hemo | 13 | EF1a-R |
| EF1a | M1-95 | Hemo | 2 | EF1a-R |
| EF1a | M1-95 | Hemo | 50 | EF1a-R |
| EF1a | M1-95 | Hemo | 58 | EF1a-R |
| EF1a | M1-95 | Hemo | 43 | EF1a-R |
| EF1a | M1-95 | Hemo | 44 | EF1a-R |
| EF1a | M1-95 | Hemo | 3 | EF1a-R1 |
| EF1a | M1-95 | Hemo | 30 | EF1a-R1 |
| EF1a | M1-95 | Hemo | 12 | EF1a-S |
| EF1a | M1-95 | Hemo | 21 | EF1a-S |
| EF1a | M1-95 | Hemo | 41 | EF1a-S |
| EF1a | M1-95 | Hemo | 42 | EF1a-S |
| EF1a | M2-22 | Foot | 9 | EF1a-R |
| EF1a | M2-22 | Foot | 107 | EF1a-R |
| EF1a | M2-22 | Foot | 130 | EF1a-R |
| EF1a | M2-22 | Foot | 138 | EF1a-R |
| EF1a | M2-22 | Foot | 14 | EF1a-R |
| EF1a | M2-22 | Foot | 36 | EF1a-R |
| EF1a | M2-22 | Foot | 64 | EF1a-R |
| EF1a | M2-22 | Foot | 65 | EF1a-R |
| EF1a | M2-22 | Foot | 66 | EF1a-R |
| EF1a | M2-22 | Foot | 15 | EF1a-R1 |
| EF1a | M2-22 | Foot | 38 | EF1a-R1 |
| EF1a | M2-22 | Foot | 1 | EF1a-S |
| EF1a | M2-22 | Foot | 4 | EF1a-S |
| EF1a | M2-22 | Foot | 19 | EF1a-S |
| EF1a | M2-22 | Foot | 20 | EF1a-S |
| EF1a | M2-22 | Foot | 21 | EF1a-S |
| EF1a | M2-22 | Hemo | 35 | MINOR |
| EF1a | M2-22 | Hemo | 1_M13 | MINOR |
| EF1a | M2-22 | Hemo | 43 | MINOR |
| EF1a | M2-22 | Hemo | 46 | MINOR |
| EF1a | M2-22 | Hemo | 2_M13 | MINOR |
| EF1a | M2-22 | Hemo | 40 | MINOR |
| EF1a | M2-22 | Hemo | 5 | MINOR |
| EF1a | M2-22 | Hemo | 60 | MINOR |
| EF1a | M2-22 | Hemo | 58 | EF1a-R |
| EF1a | M2-22 | Hemo | 3_M13 | EF1a-R |
| EF1a | M2-22 | Hemo | 6_M13 | EF1a-R |
| EF1a | M2-22 | Hemo | 3_M13 | EF1a-R |
| EF1a | M2-22 | Hemo | 56 | EF1a-R |
| EF1a | M2-22 | Hemo | 62 | EF1a-R |
| EF1a | M2-22 | Hemo | 18 | EF1a-S |
| EF1a | M2-22 | Hemo | 51 | EF1a-S |
| EF1a | M2-22 | Hemo | 57 | EF1a-S |

Table S2. NCBI genbank accession numbers of alleles found in this study

| Locus | Allele name | Origin | Accession Number |
|---|---|---|---|
| COI | U1 | Cancer | MZ724668 |
| COI | 1 | Cancer | MZ724669 |
| COI | 2 | Cancer | MZ724670 |
| COI | 4 | Host | MZ724671 |
| COI | 6 | Host | MZ724672 |
| COI | 7 | Host | MZ724673 |
| COI | 9 | Host | MZ724674 |
| COI | 10 | Host | MZ724675 |
| COI | 11 | Host | MZ724676 |
| COI | 12 | Host | MZ724677 |
| COI | 13 | Host | MZ724678 |
| COI | 14 | Host | MZ724679 |
| CR | Z1 | Cancer | MZ751085 |
| CR | 1 | Cancer | MZ751086 |
| CR | 2 | Cancer | MZ751087 |
| CR | 9 | Host | MZ751088 |
| CR | 10 | Host | MZ751089 |
| CR | 11 | Host | MZ751090 |
| EF | S | Cancer | MZ751091 |
| EF | R | Cancer | MZ751092 |
| EF | R1 | Cancer | MZ751093 |
| EF | G | Cancer | MZ751094 |
| EF | H | Cancer | MZ751095 |
| EF | H1 | Cancer | MZ751096 |
| EF | 10 | Host | MZ751097 |
| EF | 11 | Host | MZ751098 |
| EF | 12 | Host | MZ751099 |
| EF | 13 | Host | MZ751100 |
| EF | 14 | Host | MZ751101 |

Supplementary Table S3. Types of polyA region detected in cloned sequences of CR. The nucleotide sequence of polyA region in CR consists of mononucleotide sequence of adenine interrupted by one cytosine nucleotide. The sequence of polyA here is coded here as a number of adenine nucleotides before and after the cytosine.

| Individual | Tissue | Type of allele | Allele ID | 6A:C:10A | 5A:C:9A | 6A:C:9A | 6A:C:8A | 5A:C:10A | 5A:C:8A | 5A:C:11A | 6A:C:3A | 6A:C:2A | 6A:C:4A | 6A:C:6A |
|---|---|---|---|---|---|---|---|---|---|---|---|---|---|---|
| M1-105 | Foot | Healthy | CR-9 | 0 | 5 | 0 | 0 | 13 | 0 | 1 | 0 | 0 | 0 | 0 |
| M1-105 | Hemo | Cancer | CR-2 | 16 | 0 | 0 | 0 | 0 | 0 | 0 | 0 | 0 | 0 | 0 |
| M1-162 | Foot | Cancer | CR-1 | 0 | 5 | 0 | 0 | 0 | 0 | 0 | 0 | 0 | 0 | 0 |
| M1-162 | Foot | Healthy | CR-10 | 12 | 0 | 0 | 0 | 0 | 0 | 0 | 0 | 0 | 0 | 0 |
| M1-162 | Hemo | Cancer | CR-1 | 0 | 17 | 0 | 0 | 4 | 3 | 0 | 0 | 0 | 0 | 0 |
| M1-95 | Foot | Cancer | CR-Z1 | 1 | 0 | 1 | 0 | 0 | 0 | 0 | 0 | 0 | 0 | 0 |
| M1-95 | Foot | Healthy | CR-11 | 0 | 0 | 0 | 0 | 0 | 0 | 0 | 5 | 1 | 3 | 1 |
| M1-95 | Hemo | Cancer | CR-Z1 | 1 | 0 | 26 | 0 | 0 | 0 | 0 | 0 | 0 | 0 | 0 |
| M2-22 | Foot | Cancer | CR-Z1 | 0 | 0 | 11 | 1 | 0 | 0 | 0 | 0 | 0 | 0 | 0 |
| M2-22 | Hemo | Cancer | CR-Z1 | 0 | 0 | 17 | 0 | 0 | 0 | 0 | 0 | 0 | 0 | 0 |
